# Fluorogenic speed-optimized DNA-PAINT probes enable super-resolution imaging of whole cells

**DOI:** 10.64898/2026.03.23.710523

**Authors:** Sylvi Stoller, Asmita Jha, Joerg Bewersdorf, Florian Schueder

**Author notes:** Corresponding authors. (J.B.) & (F.S.). These authors contributed equally (Sylvi Stoller, Asmita Jha).

## Abstract

Super-resolution microscopy with DNA-PAINT enables molecular-scale, multiplexed, and quantitative imaging, but its throughput is limited by slow binding kinetics and elevated background at high probe concentrations. Recent speed-optimized and fluorogenic probes improve performance but impose strong constraints on sequence design, revealing a fundamental tradeoff between fast binding and efficient quenching. Here, we introduce a modular probe architecture that spatially decouples binding kinetics from fluorophore–quencher interactions by integrating speed-optimized sequence motifs with PEG spacers. Using DNA origami nanostructures, we demonstrate enhanced localization rates, signal-to-background ratios, and imaging efficiency compared to state-of-the-art probes. We validate our approach in cells, demonstrating its capability to image nuclear targets and enabling three-dimensional imaging of the endoplasmic reticulum using standard widefield illumination. Our work establishes a general framework for fast, multiplexed, and low-background super-resolution imaging.

## Introduction

The development of super-resolution microscopy has revolutionized the field of biology, allowing scientists to investigate molecular organization well below the diffraction limit of light^1,2^. In single-molecule localization microscopy (SMLM), this is achieved by having only a sparse subset of labeled target molecules emitting fluorescence (i.e. ‘blinking’) at any given time. After capturing these spatially-isolated blinks across thousands of frames, their positions can be individually localized with sub-diffraction precision, resulting in approximately 20-30 nm lateral and 50-75 nm axial resolution^3–8^.

DNA points accumulation for imaging in nanoscale topography (DNA-PAINT)^9^ is an implementation of SMLM that generates single-molecule blinks through short, fluorescently-labeled, single-stranded DNA (ssDNA) ‘imager probes’ transiently binding to complementary ssDNA ‘docking strands’, which are attached to the targets of interest. While unbound, these probes diffuse freely in solution, appearing undetectable to the camera but contributing to a homogenous background. When probes are bound to a docking strand, they become immobilized and produce a bright fluorescent signal that the camera detects as a blinking event. The binding kinetics of the probe can be modified by the DNA sequence, making DNA-PAINT a highly programmable, versatile, and robust super-resolution technique.

The spatial resolution of DNA-PAINT depends on the precise localization of single molecules, which is limited by the brightness of individual blinking events relative to the diffuse background generated by unbound probes. High localization precision is typically achieved by (1) reducing the probe concentration to limit the background level or (2) restricting the excitation or detection volume through optical sectioning techniques including total internal reflection fluorescence (TIRF) microscopy^10^, highly inclined and laminated optical sheet (HILO) illumination^11,12^, confocal microscopy^13,14^ or light-sheet microscopy^15,16^. However, the use of low probe concentration reduces the imaging speed while optical sectioning increases microscope hardware complexity and restricts biological application.

In recent years, there have been two major approaches to developing specialized DNA-PAINT probes that help overcome these limitations. The first is to decrease the detected fluorescence from unbound probes, allowing the use of higher probe concentrations without increasing background levels. FRET DNA-PAINT relies on Förster Resonance Energy Transfer (FRET) to create a strong signal only when a donor-labeled probe is bound to a docking site with an acceptor^17,18^ Fluorogenic DNA-PAINT creates self-quenching probes by conjugating a dye-quencher pair at opposite ends of the probe sequence, leading to a ∼50-fold increase in fluorescence upon binding^19^. The second approach is to increase the association rate (k_on_) of the probe, providing high blinking rates even at low probe concentrations. Ago-PAINT, for example, which loads imager probe strands into Argonaute proteins, allows 10-fold faster target binding^19^, and Speed-optimized DNA-PAINT, which leverages a short, speed-optimized sequences and repetitive docking sites, achieves up to a 100-fold increase in target sampling over previous DNA-PAINT probes^20,21^.

While both approaches have been successful, these developments have thus far been mutually exclusive tools, requiring users to make a trade-off between imaging speed and background level. Among these, Fluorogenic DNA-PAINT and Speed-optimized DNA-PAINT are the only tools that rely on a single, modified-DNA probe, making them the most suitable candidates for integration. However, the former requires a long (15 nt) DNA strand to sufficiently separate the dye and quencher for strong fluorogenic effects from FRET quenching, while the latter relies on short (6-7 nt), specialized probe sequences designed for high association rates.

Here we introduce a new class of DNA-PAINT probes, which we term fluorogenic speed-optimized probes (FSPs), that overcome this fundamental contradiction. We combine the self-quenching concept of Fluorogenic DNA-PAINT with the short sequence design from Speed-optimized DNA-PAINT by inserting inert polyethylene glycol (PEG) spacers at both ends of the DNA strand to create sufficient distance between the dye and quencher. FSPs exhibit low diffuse background in their unbound state while simultaneously featuring high association rates even under non-TIRF illumination, enabling fast, widefield-based super-resolution imaging of whole cells without optical sectioning.

## Results

### Fluorogenic speed-optimized probe (FSP) design

High fluorogenicity, high association rates, and low unspecific binding are key considerations when selecting a DNA-PAINT probe. Fluorogenic DNA-PAINT achieves fluorogenicity with self-quenching probes that have a fluorophore and quencher conjugated at opposing ends of the DNA sequence^17^. In the unbound, single-stranded state, the short persistence length of DNA (∼1 nm)^22,23^ facilitates interaction between the fluorophore and quencher. Upon binding to the docking strand, the rigid DNA duplex^24^ spatially separates the fluorophore and quencher, reducing interaction and allowing fluorescence (**Fig. 1a**, top left). To minimize the quenching effect in the bound state, Fluorogenic DNA-PAINT probes are 15 nt long so that their duplex form exceeds the Förster radius^25^. However, longer DNA sequences are more prone to secondary structures that can lead to slow association rates, heightened crosstalk, and increased non-specific binding to cellular components^26^.

**Figure 1.**
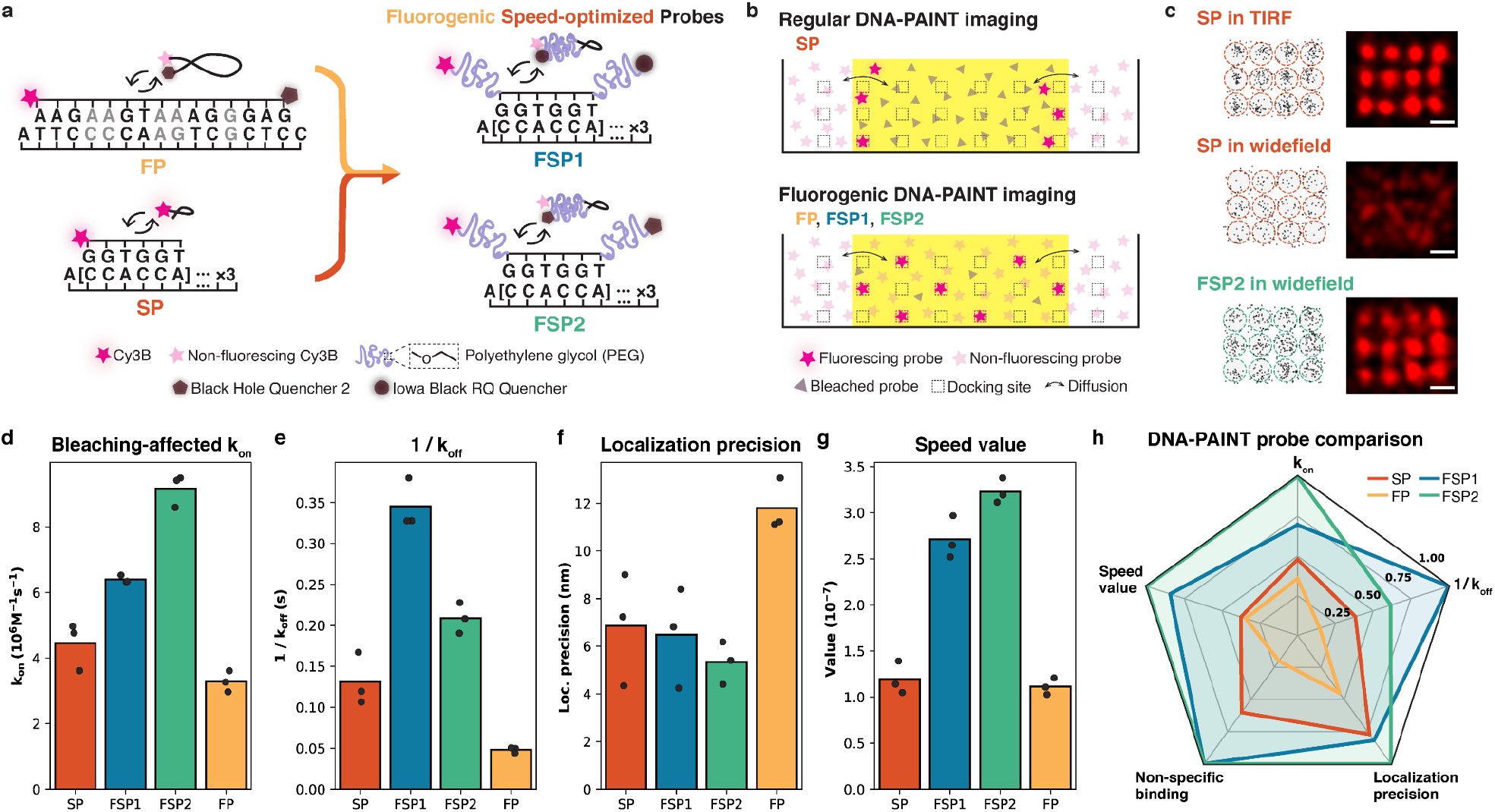
Fluorogenic speed-optimized probe (FSP) design and characterization. (a) FSPs (FSP1 and FSP2) integrate probe and docking site sequences from Speed-optimized DNA-PAINT probe SP with the fluorophore-quencher design from Fluorogenic DNA-PAINT probe FP. Grey letters indicate locations of single-nucleotide mismatches in FP. (b) Bleaching effect in DNA-PAINT imaging under widefield illumination (yellow area) with regular and fluorogenic probes. (c) DNA-PAINT imaging of 20-nm grid DNA origami demonstrating that SP loses its ability to resolve the grid structures going from TIRF to widefield, but FSPs can resolve the structure under widefield illumination (**Suppl. Fig. 3b**). Scale bars: 20 nm. (d-g) Comparison of binding kinetics (k_on_ and 1/k_off_), localization precision, and speed value measured on DNA origami (n=3 independent experiments for each probe, each value averaged over ∼2000 origami) (see **Suppl. Fig. 3d-h** for more). (h) Overall DNA-PAINT probe comparison.

Speed-optimized DNA-PAINT, on the other hand, relies on short sequence motifs to achieve its high association rates^20,21,27^. The combination of repetitive 2- or 3-nt sequence motifs used in short probes (6-7 nt) with repeats of the complementary binding motif provides multiple binding sites at the docking strand (**Fig. 1a**, bottom left). Additionally, the sequence of each motif is limited to two letters to preclude secondary structure formation, ensuring that all probes in solution can readily bind to the docking strand.

To reconcile the fast association rates of Speed-optimized DNA-PAINT with the fluorogenic properties of Fluorogenic DNA-PAINT, we integrated a fluorophore-quencher pair with an established short, speed-optimized sequence by inserting PEG spacers on both ends of the DNA sequence (**Fig. 1a**, right). PEG is a synthetic polymer formed from repeating ethylene glycol units which is known to be water-soluble and chemically inert, making it an ideal candidate for this purpose^28^. We chose PEG spacers with nine ethylene glycol units (PEG9), which result in a total estimated contour length of ∼5 nm^29,30^. This hybrid probe architecture conceptually decouples the hybridization kinetics from the fluorophore-quencher interactions, and we hypothesized that this design would yield a probe with both a high association rate and fluorogenicity.

For the DNA sequence, we chose the R2 sequence since it has the highest association rate of the original six Speed-optimized DNA-PAINT probes^27^. As a fluorophore, we use Cy3B, one of the brightest and best-established dyes for DNA-PAINT^31^. In addition to the Black Hole Quencher 2 (BHQ2), which was paired with Cy3B in the original Fluorogenic DNA-PAINT implementation^17^, we introduce another probe with Iowa Black Red Quencher (IBRQ) because of its favorable spectral overlap with the Cy3B emission spectrum and stabilizing effect on DNA duplexes^24^. Our two new probes are termed fluorogenic speed-optimized probes 1 and 2 (FSP1 and FSP2) (**Fig. 1a**, right).

In addition to the anticipated background reduction, we hypothesized that the FSPs would also have increased resistance to bleaching, thereby facilitating imaging in non-TIRF illumination regimes. With typical DNA-PAINT imaging, in large excitation volumes, the replenishment of probes from solution becomes less efficient. In widefield illumination, for example, probes must diffuse over long distances and may bleach in solution before ever reaching the intended target (**Fig. 1b**, top). This results in a reduced number of observed blinks, resembling a lower effective association rate. Since photobleaching is associated with the excited state of the fluorophore, we reasoned that a probe will be resistant to bleaching while quenched. Therefore, fluorogenic probes like FSPs should provide a more uniform availability of unbleached probes throughout the illuminated volume (**Fig. 1b**, bottom) (**Suppl. Note 1**), making them uniquely suited for volumetric imaging for biological applications.

For benchmarking, in the following experiments, we compared the new FSPs with the original Speed-optimized DNA-PAINT probe R2^27^, here called ‘SP’, and a Fluorogenic DNA-PAINT probe with the BHQ2-Cy3B pair and the sequence of Imager probe B, which has the higher association rate of the two original sequences^17,32^, here called ‘FP’. This way, all probes share the same dye and FSP2 and FP share the same quencher, while FSP1, FSP2, and SP share the same binding motif.

### Single-molecule characterization of DNA-PAINT probes

We first tested the performance of our new FSPs by imaging DNA origami with 12 docking sites arranged in a 20-nm grid^33,34^ (**Suppl. Fig. 3a**) and comparing them to SP^27^. While SP excels in the TIRF regime and could resolve the grid structure (**Fig. 1c**, top), it was unable to resolve the structure under widefield illumination (**Fig. 1c**, middle) due to the increased background. In contrast, FSPs were able to clearly resolve the 20-nm grid structure even under widefield illumination (**Fig. 1c**, bottom; **Suppl. Fig. 3b**).

To compare FSP1 and FSP2 more quantitatively to the original SP^27^ and FP^17,32^, we measured their binding kinetics under non-TIRF illumination using DNA origami structures featuring a single docking site, as used in previous publications^35,36^ (**Suppl. Fig. 3c**). The most informative metrics were the association rate (k_on_), the ON time (i.e. the inverse of the effective dissociation rate k_off_), the localization precision, and the ‘speed value’, a metric describing the number of blinks relative to the background level (**Suppl. Note 1**). SP and FP were imaged at laser intensities of 0.2 and 4.1 kW/cm^2^, respectively, close to those reported in previous publications^17,22^. For the FSPs, we measured kinetics across laser intensities ranging from ∼0.2 to ∼7 kW/cm^2^ (**Suppl. Fig. 2**) and selected ∼1.5 kW/cm^2^ for FSP1 and ∼4.1 kW/cm^2^ for FSP2 as the optimal intensities based on the binding kinetics and photophysical properties of these probes (**Suppl. Note 2**).

Under these conditions, we found FSP1 and FSP2 to have association rates of 6.4±1.5 and 9.2±2.2 *10^6^/(M*sec) (mean±s.e.m.), respectively, ∼2 to 3-fold higher than FP (3.3±0.6 *10^6^/(M*sec)) and ∼1.3 to 2-fold higher than the effective association rate of SP (4.4±1.2*10^6^/M*sec)) (**Fig. 1d**). We note that the true association rate of SP is likely higher, since our measured rate is reduced by bleaching caused by widefield illumination (**Suppl. Note 1**)^22^

For the ON time, 1/k_off_, we measured 0.34±0.07 s and 0.21±0.04 s for FSP1 and FSP2, respectively (**Fig. 1e**). These values are higher than that of FP (0.05±0.02 s), as FP lacks the k_on_ optimization of SP. Additionally, they are elevated compared to SP’s ON time (0.13±0.04 s), which we hypothesize is reduced due to decreased blinking detection efficiency and increased bleaching of SP (**Suppl. Note 1**).

FSP1 and FSP2 have the best localization precision of the four probes (**Fig. 1f**), achieving 5.3±1.0 nm and 6.5±1.8 nm, respectively, compared to 6.9±2.4 nm for SP and 11.8±2.2 nm for FP.

In DNA-PAINT, the background level, and therefore the localization precision and SBR, depend on the probe concentration. Reducing the concentration leads to better localization precision; however, this comes at the cost of fewer localization events per second, resulting in longer imaging times. To capture this tradeoff between sampling rate and background contribution of DNA-PAINT probes, we introduce the ‘speed value’, a concentration-independent metric defined as the ratio of binding events per second to the number of background photons per second (**Suppl. Note 1**). Probes that generate many blinks while maintaining low background have a high speed value. FSP1 and FSP2 yield the highest speed values of 2.7±1.2 and 3.2±1.4 (*10^-7^), respectively, compared to 1.1±0.4 and 1.2±0.5 (*10^-7^) for SP and FP, respectively (**Fig. 1g**).

We provide a qualitative comparison of the single-molecule characterizations of the four probes complemented by an assessment of non-specific binding in biological samples (**Fig. 1h**) by representing each property such that the highest value is most desirable (see Methods).

FSP1 and FSP2 have the largest total area, which speaks to the superior performance of our new probes over the current gold standards.

### Benchmarking of FSPs for cellular imaging

Next, we confirmed that FSPs are also capable of high-quality super-resolution imaging under widefield illumination in a cellular environment by imaging microtubules in COS-7 cells. We immunolabeled alpha-tubulin with a transient adapter docking site (**Suppl. Table 2**), recently introduced in FLASH-PAINT^32^, and imaged the same field of view with both FSP1 and FSP2 (**Fig. 2a, Suppl. Fig. 2**), achieving 9.63 nm and 6.91 nm NeNA localization precision^36^, respectively. Both FSPs readily resolve the labeled microtubules as hollow filaments with a Fourier ring correlation (FRC) resolution^35^ of 33 nm for FSP1 and 28 nm for FSP2 (**Fig. 2b**). We measured the effective diameter of the highlighted microtubule segment to be 51 nm and 46 nm for FSP1 and FSP2, respectively (**Fig. 2c**), consistent with the expected width of antibody-coated microtubules^37,38^. As expected from the kinetics, FSP2 achieves a slightly better localization precision and image resolution than FSP1.

**Figure 2.**
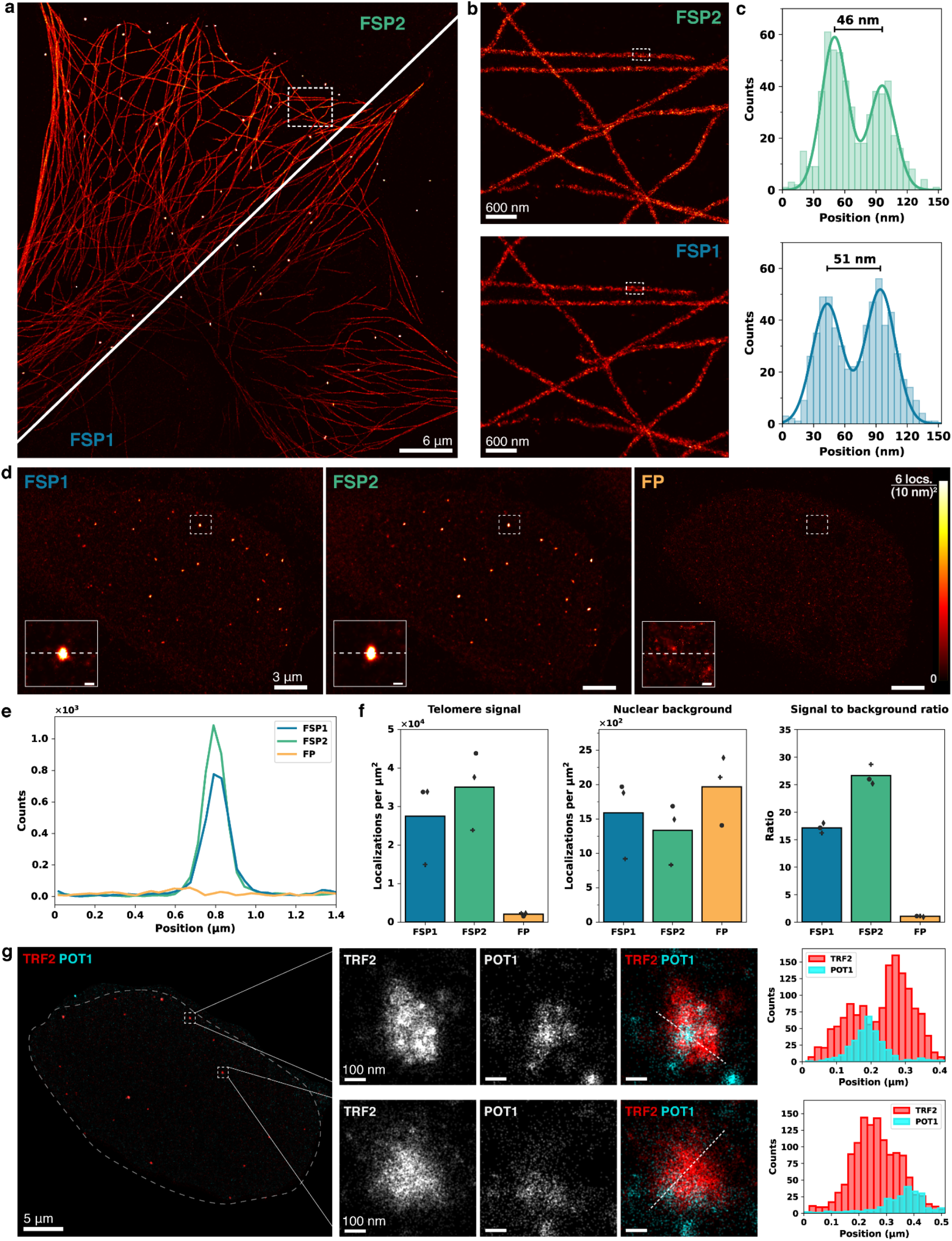
Validation of FSPs for cellular and nuclear imaging. (a) DNA-PAINT image of microtubules in a fixed COS-7 cell imaged with FSP2 (top left) and FSP1 (bottom right) under widefield illumination. (b) Zoomed-in image of boxed region in a for each probe. (c) Cross-sectional histograms of the boxed microtubule segment in b with double Gaussian fit to obtain peak-to-peak distances. (d) Three rounds of DNA-PAINT imaging of telomeres in a U-2 OS cell with FSP1, FSP2, and FP. Insets highlight the same region in each nucleus; FP inset displayed at 2x intensity (scale bar: 200 nm). (e) Line profiles along the white line shown in each inset in d. (f) Comparison of telomere signal (left), nuclear background (middle), and signal-to-background ratio (right) between the three fluorogenic DNA-PAINT probes. Each marker shape represents data collected from an independent experiment (n=3) (**Suppl. Fig. 4**). (g) Multiplexed FLASH-PAINT image of U-2 OS cell nucleus (outlined) with overexpressed Myc-POT1 and endogenous TRF2 targeted. Dashed line indicates approximate boundary of segmented nucleus; localizations outside are not displayed (see Methods). Zoom-ins highlight the distinct spatial distribution of TRF2 and POT1 within clusters; Pot1 displayed at 2x intensity. The histograms were taken along the white lines shown in the neighboring insets.

### High signal-to-background ratio with FSPs in the nucleus

We next tested our FSPs’ compatibility with nuclear imaging, which has historically been challenging for DNA-PAINT due to increased non-specific binding and impeded diffusion in the nucleus^39,40^. Longer DNA sequences in particular are more susceptible to binding non-specifically to endogenous RNAs or DNAs, so the long FPs therefore tend to exhibit increased background in nuclear environments. Due to the short DNA sequence of FSPs and their robust performance under widefield illumination, we expected the FSPs to be uniquely suited for imaging within the nucleus. First, we imaged telomeres labeled with DNA fluorescence in-situ hybridization (DNA-FISH) in U-2 OS cells. To allow for a direct and unbiased comparison with FP, we designed primary probes complementary to telomeric repeats with an FSP docking site on the 3’ end and an FP docking site on the 5’ end. We performed sequential rounds of DNA-PAINT imaging via Exchange-PAINT^41^ on the same nucleus with the FSPs and FP (**Fig. 2d, Suppl. Fig. 4**). While telomere puncta are easily identifiable for both the FSP1 and FSP2 images, there is no clear signal seen in the FP round. Along with the loss in signal, we see a ∼20-50% increase in nuclear background caused by non-specific binding of FP (**Fig. 2f**, middle). Combined, FSPs yield a significantly higher signal-to-background ratio compared to FP (∼16-fold and ∼25-fold, respectively) (**Fig. 2f**, right).

DNA-PAINT traditionally excels in imaging abundant cellular targets where the probe concentration can be kept low. To further test FSP performance in the nucleus for targets less abundant than telomeric repeats, we imaged the shelterin complex, which forms a cap on the ends of telomeres and protects them from degradation and DNA damage. We selected two subunits of the shelterin complex^42^: telomere repeat factor 2 (TRF2), which binds double-stranded telomeric DNA, and protection of telomeres 1 (POT1), which binds single-stranded telomeric DNA. We labeled overexpressed Myc-POT1^43^ and endogenous TRF2 in U-2 OS cells with primary antibodies and DNA-conjugated secondary nanobodies with FLASH-PAINT transient adapter^32^ sequences. Sequential imaging of the two targets revealed that the two proteins form puncta throughout the nucleus (**Fig. 2g**, left)^44^. POT1 and TRF2 puncta appear in close vicinity to each other but display distinct spatial distributions within each cluster (**Fig. 2g**, zoom-ins), consistent with POT1 and TRF2 being part of two different subcomplexes that bind distinct regions of the telomeric chromatin^42,44^. The TRF2 puncta generally exhibit higher signal levels than POT1 (**Fig. 2g**, right), consistent with its higher relative abundance^42^. These results demonstrate that FSPs extend the imaging capabilities of DNA-PAINT to previously elusive, low-abundance targets in challenging imaging environments such as the crowded cell nucleus.

### Whole-cell 3D imaging of the endoplasmic reticulum without optical sectioning

We next focused our attention on applications that are a hallmark of cell biological imaging, but among the most challenging use cases for super-resolution microscopy: volumetric, whole-cell imaging of densely labeled structures. Achieving a holistic view of the cell requires imaging not just at the coverslip surface or in a single slice but across depths that span several micrometers. We imaged the endoplasmic reticulum (ER) in a U-2 OS cell overexpressing mEmerald-Sec61β using an anti-GFP nanobody conjugated to a FSP docking site (**Suppl. Table 2**). We collected a series of Z-stacks over a depth of ∼6 µm with a 300 nm step size and acquired a total of ∼200 million localizations with 3.3 nm average localization precision and 14 nm NeNA localization precision^36^. **Fig. 3a-g** show that we are able to resolve the complete ER tubular network, including single ER tubules at the periphery of the cell as well as the densely packed perinuclear ER. **Fig. 3i and 3j** in particular highlight contact sites between ER tubules and the outer membrane of the nuclear envelope.

**Figure 3.**
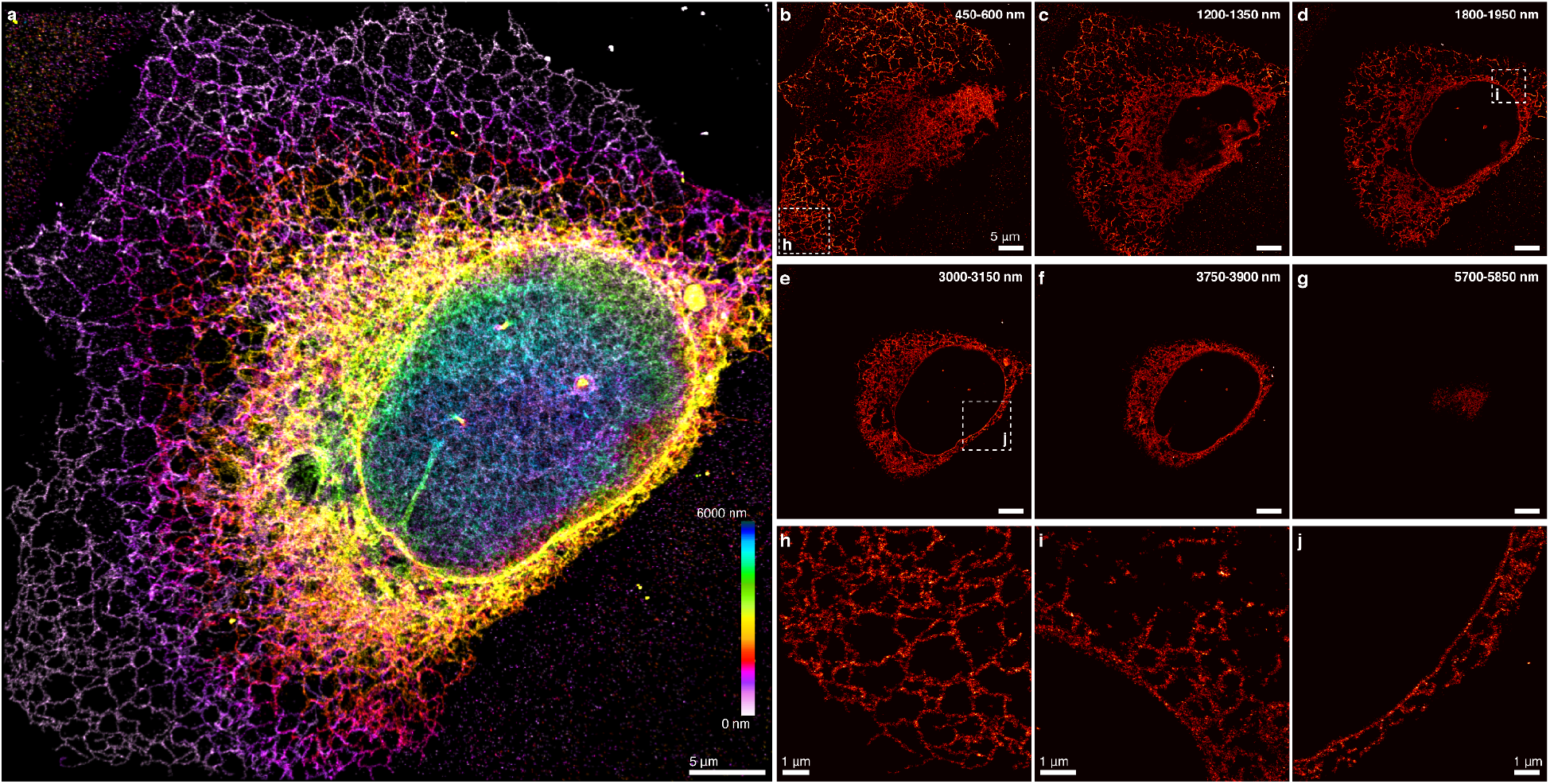
Whole-cell 3D DNA-PAINT imaging of the endoplasmic reticulum without optical sectioning. (a) 3D DNA-PAINT imaging of the endoplasmic reticulon (ER) with FSP2. (b-g) Representative 150-nm-thick slices spread throughout the full height of the cell, from the layer of ER network between the nuclear envelope and plasma membrane (b) to the ER network at the top of the cell (g). (h-j) Zoomed-in, 75-nm-thick views of the boxed regions in b, d, and e, showing the well-resolved intricate network of the ER at the cell periphery as well as in the dense perinuclear space, including connections between the ER and nuclear envelope. Note that ER tubules can appear fragmented due to the thin Z-sections displayed.

## Discussion

With FSPs, we have demonstrated the first integration of chemical spacers into DNA-PAINT probes. This conceptual advance combines two previously incompatible strengths, the high association rate from Speed-optimized DNA-PAINT^20,21,27^ and the fluorogenicity of Fluorogenic DNA-PAINT^17^, into a single probe. This integration has enabled fast super-resolution imaging under widefield illumination, eliminating the need for optical sectioning.

Suppressing background from non-fluorogenic probes has thus far relied on optical sectioning techniques to reduce out-of-focus signal. These techniques, including TIRF, HILO, light-sheet microscopy and confocal microscopy variants, all require relatively complex optics compared to standard widefield illumination. FSPs eliminate this need and are, in principle, even compatible with strong LED light sources making lasers obsolete.

In addition to alleviating instrumentation requirements, FSPs open new possibilities for studying nuclear biology. Imaging nuclear targets with DNA-PAINT has been a longstanding challenge due to the dense nuclear environment. Previously, performing DNA-PAINT in the nucleus has relied on specialized, costly probes based on left-handed DNA^39^ or a combination of physical and optical sectioning^16,45,46^. By contrast, FSPs directly provide a high signal-to-background ratio in the nucleus at a reagent cost comparable to Fluorogenic DNA-PAINT. For applications where non-specific antibody binding may be a substantial source of background, FSPs are conceptually compatible with approaches such as Shielded DNA-PAINT which have been designed to mitigate labeling-related issues^40^.

The FSPs have low non-specific binding and are fluorogenic, which protects them from premature bleaching while they travel to their target sites. These features make FSPs an ideal choice for complex biological samples, where high background and slow diffusion across large distances are known challenges.We expect FSPs to be uniquely suited for super-resolution imaging of multicellular systems such as 3D cell cultures, organoids, tissues, or small model organisms.

The modular design of the FSPs allows for the straightforward extension of the FSP toolbox to orthogonal DNA sequences and spectrally distinct dyes. Furthermore, we have shown in this work that FSPs are compatible with the transient adapters of FLASH-PAINT^32^ and expect that they will work equally well with other multiplexing strategies such as SUM-PAINT^47^. Additionally, we anticipate that the fluorogenicity and reduced non-specific binding of FSPs will benefit MINFLUX^47,48^and RESI^49^, recent SMLM developments capable of achieving ∼1 nm resolution.

Beyond their immediate impact in super-resolution microscopy, with strengths in volumetric imaging and multiplexing capabilities, FSPs have the potential to enable DNA-technology based applications in spatial transcriptomics, chromatin tracing, and even diffraction-limited, gentle multiplexed imaging at the tissue level.

## Acknowledgements

We thank Felix Rivera-Molina for support and advice regarding imaging on the Dragonfly microscope and the CINEMA lab facility at Yale for enabling the use of the microscope for our imaging experiments. We thank Sandy Chang for sharing the POT1 plasmid with us. We thank Kenny Chung for fruitful discussions and advice regarding the concept of fluorogenic imager probes. A.J. is supported by an NIH F31 fellowship (AR085501) and an NIH T32 (GM145469-03). S.S. is supported by Yale’s Integrated Graduate Program in Physical and Engineering Biology. J.B. and S.S. acknowledge support from the National Institutes of Health (GM151829) and the National Science Foundation (2520354). F.S. acknowledges support from the Human Frontier Science Program (LT000056/2020-C) and funding from EPFL to the Laboratory of Molecular Spatial Omics.

## Author contributions

F.S. conceived the concept of the new probe. S.S. and A.J. performed the experiments. A.J. prepared the cell samples. S.S. and A.J. carried out the cell imaging. S.S. prepared and imaged the DNA origami samples and performed data analysis. F.S. and J.B. supervised the study. All authors contributed to interpreting the results, writing, and editing the manuscript.

## Data availability statement

All raw data is available upon reasonable request from the authors.

## Code availability statement

Picasso^33^, used for data analysis and visualization, can be downloaded from Github (https://github.com/jungmannlab/picasso). PYMEVisualize^50^, used for visualization, can be downloaded from Github (https://github.com/python-microscopy/python-microscopy).

## Materials and Methods

### Buffers

Four buffers were used for DNA origami sample preparation and imaging. Buffer A (10 mM Tris pH 8.0 (invitrogen, AM9856), 100 mM NaCl (invitrogen, AM9759)); Buffer A+ (Buffer A supplemented with 0.05% Tween 20 (MilliporeSigma, P9416-50mL)); Buffer B+ (10 mM MgCl_2_ (invitrogen, AM9530G), 5 mM Tris-HCl pH 8.0 (invitrogen, AM9856), 1 mM EDTA pH 8.0 (invitrogen, AM9261), 0.05% Tween 20); Buffer C (1× PBS (Gibco), 500 mM NaCl (invitrogen, AM9759)). For experiments employing an oxygen scavenging system (as indicated in **Suppl. Table 3**), the imaging buffer was supplemented with Trolox (Santa Cruz Biotechnology, sc-200810A) dissolved in DMSO (invitrogen, D12345) that was stored at -20 °C in 50mM aliquots, and sodium sulfite (MilliporeSigma, S0505).

### DNA origami self-assembly

All DNA origami structures were designed with the Picasso design tool.^33^ Self-assembly of DNA origami was accomplished in a one-pot reaction with 50 μL total volume, consisting of 10 nM M13mp18 scaffold strand (New England BioLabs, N4040S), 10 nM biotinylated staples (IDT) (**Suppl. Table 4**), and 100 nM folding staples (IDT) and 1 μM of docking site strands (**Suppl. Tables 5** and **6**) in folding buffer (125 mM MgCl_2_, 100 mM Tris pH 8.0 and 10 mM EDTA pH 8.0). The reaction mix was then subjected to a thermal annealing ramp using a thermocycler. The reaction mix was first incubated at 80 °C for 5 min, then cooled from 60 to 4 °C in steps of 1 °C every 3.21 min, and then held at 4 °C.

### DNA origami sample preparation

For DNA origami sample preparation, a µ-Slide VI^0.5^ (ibidi, 80607) was used as a sample chamber. First, 100 μL of biotin-labeled bovine albumin (MilliporeSigma, A8549) at 1mg/mL in Buffer A+ (stored at -20 °C at 10mg/mL in Buffer A) was flushed into the chamber and incubated for 5 min. Then, the chamber was washed with 500 μL of buffer A+ and 100 μL of streptavidin (ThermoFisher, S-888) at 0.5 mg/mL in buffer A) was flushed into the chamber and allowed to bind for 5 min. After washing with 500 μL of buffer A and subsequently with 500 μL of buffer B, 100 μL of biotin-labeled DNA origami structures (∼100-300 pM depending on desired density) in buffer B were flushed into the chamber and incubated for 10 min. Finally, after DNA origami incubation, the chamber was washed with 500 μL of buffer B.

### Cell culture and transfection

U-2 OS (ATCC, HTB-96 Lot# 70008732) and COS-7 (ATCC, Lot #63624240) were maintained in DMEM, high glucose (Gibco, 11965092) and 10% FBS (Gibco, A5256801) in a 5% CO2 environment. Subculturing was performed using 0.25% Trypsin (Gibco, 25200056) and 1× PBS (Gibco, 10010023).

### Cell sample preparation

Cells were seeded at 70-80% confluency on 8-well glass bottom slides (Ibidi, 80827) and then processed as described below. All incubations for samples were done at room temperature with sample rocking, unless otherwise noted.

#### Microtubule sample preparation

Microtubules were labeled following a previously published protocol^51^. COS-7 cells were incubated with 0.2% saponin (MilliporeSigma, SAE0073) for 45 seconds, and then immediately fixed with 3% formaldehyde (Electron Microscopy Sciences, 15710) and 0.1% glutaraldehyde (Electron Microscopy Sciences, 16019) in 1× PBS for 15 minutes, rinsed 3 times with 1× PBS (Gibco, 10010023), and incubated in blocking buffer (1× PBS + 0.2% Triton X-100 (MilliporeSigma, T8787) + 3% bovine serum albumin (Millipore Sigma, A8549) for 30 min. Samples were then incubated in mouse anti-alpha tubulin primary antibody (MilliporeSigma, T5168) at a concentration of 1:250 in antibody dilution buffer (1× PBS + 1% BSA + 0.2% TX-100) overnight at 4 °C. The cells were then washed 3 times for 5 minutes each with wash buffer (1× PBS + 1% BSA + 0.05% Triton X-100) and incubated with oligonucleotide-conjugated goat anti-mouse IgG secondary antibody (Jackson ImmunoResearch, 115-005-146) at a concentration of 1:250 for 1 hour at room temperature. The secondary antibody was conjugated to FLASH-PAINT adapter oligonucleotide docking strands using azide/DBCO click chemistry^32^. Finally, cells were post-fixed with 3% formaldehyde + 0.1% glutaraldehyde in 1× PBS for 10 minutes, washed 3 times with 1× PBS, and imaged immediately after. Gold nanoparticles (Microspheres Nanospheres, 790118-010) were added at 1:100 and allowed to settle before imaging.

#### Telomere sample preparation

U-2 OS cells were fixed with 4% formaldehyde in 1× PBS for 15 minutes. Samples were then washed 3 times with 1× PBS and permeabilized with room temperature permeabilization buffer (0.25% Triton X-100 in 1× PBS) for 15 minutes. They were then incubated in 0.1 M HCl (MilliporeSigma, 2104) for 5 minutes and washed 3 times with 1× PBS. Samples were then incubated in hybridization solution 1 (2× SSC (Thermo Fisher Scientific, AM9770), 50% vol/vol Formamide (Thermofisher Scientific, AM9342), and 0.1% Tween (MilliporeSigma, P7949-500mL)) for 35 minutes. The hybridization solution was then aspirated, and 175 µL of hybridization buffer 2 (2× SSC, 50% vol/vol Formamide, 0.1% Tween, 10% Dextran vol/vol (MilliporeSigma, S4030) and 2 µM telomere probe (Suppl. Table 2) were added to the samples. The dish was then placed on a 90 °C heat block for 3 minutes and then placed in a humid chamber overnight at 42 °C. The following day, samples were washed 2 times with warm 2× SSC for 10 minutes each. Samples were then post-fixed with 3% formaldehyde + 0.1% glutaraldehyde in 1× PBS for 10 minutes and washed 3 times with 1× PBS.

#### Myc-POT1 and TRF2 sample preparation

U-2 OS cells were transfected with 750 ng of POT1 expression plasmid with a Myc tag (gifted by Sandy Chang)^43^, using Lipofectamine 3000 (ThermoFisher Scientific, L3000015) 24 hours after being seeded. U-2 OS cells were then incubated for 16-24 hours and then placed in 800 µg/mL Puromycin (ThermoFisher Scientific, A1113802) selection media for 48 hours before proceeding with sample preparation. After selection, cells were fixed with 4 % formaldehyde in 1× PBS for 15 minutes. The fixative was then removed and quenched with 0.1 % sodium borohydride (Millipore Sigma, 213462-25G) in 1× PBS for 7 minutes, followed by 100 mM glycine (US Biological, 56-40-6) in 1× PBS for 10 minutes. Then samples were rinsed 3 times with 1× PBS and permeabilized with room temperature permeabilization buffer (0.25 % Triton X-100 in 1× PBS) for 15 minutes. Cells were then blocked for 1 hour with blocking buffer (1% BSA in 0.1% Triton X-100), blocking buffer was then removed, and samples were labeled at 1:300 with c-myc monoclonal antibody (ThermoFisher Scientific; 9E10 cMyc) against POT1 and TRF2 (Novus Biologicals; 57130) at 4 ºC overnight. The next day, samples were washed 3 times with 1× PBS for 5 minutes each and incubated in DNA-labeled secondary nanobodies (Massive Photonics) corresponding to their host target for 1 h in the blocking buffer^32^. The samples were then washed 3× with 1× PBS for 5 minutes each and post-fixed with 3% formaldehyde and 0.1% glutaraldehyde in 1× PBS for 10 minutes and washed 3 times with 1× PBS.

#### Sec61β sample preparation

U-2 OS cells were transfected with 1 µg of mEmerald-Sec61β, (Addgene, 54249), using Lipofectamine™ LTX Reagent with PLUS™ Reagent (ThermoFisher Scientific, 15338100) 24 hours after being seeded. Cells were incubated for 16-24 hours until they reached desirable transient expression levels before proceeding with sample preparation. Cells were fixed with 3% formaldehyde and 0.1% glutaraldehyde in 1× PBS for 15 minutes. The fixative was then removed and the sample was quenched with 0.1% sodium borohydride in 1× PBS for 7 minutes, followed by 100 mM glycine in 1× PBS for 10 minutes. Then, samples were rinsed 3 times with 1× PBS and permeabilized with room temperature permeabilization buffer for 15 minutes. Cells were then blocked for 1 hour with 1% BSA in 0.1% Triton X-100. After blocking buffer was removed, the samples were labeled at 1:300 with a GFP nanobody conjugated with a DNA-PAINT docking site (Massive Photonics) at 4 ºC overnight^33^. Samples were then washed 3 times with 1× PBS for 5 minutes each and, finally, post-fixed with 3% formaldehyde and 0.1% glutaraldehyde for 10 minutes and washed 3 times with 1× PBS. Gold nanoparticles were added at 1:100 and allowed to settle before imaging.

#### Super-resolution microscope setup

Fluorescence imaging was carried out on an inverted Nikon Eclipse Ti2 microscope (Nikon Instruments) with a Perfect Focus System, equipped with an Andor Dragonfly unit. The Dragonfly was used in the BTIRF mode, applying an objective-type TIRF or widefield configuration with an oil-immersion objective (Nikon Instruments, Apo SR TIRF 60×, NA 1.49, Oil). For TIRF imaging (**Fig. 1c**), the adjustment was set above the critical angle; for all other experiments, the adjustment was set to range from smaller than the critical angle to parallel to the optical axis of the objective lens. For excitation, a 561-nm laser (1 W nominal laser power) was used. The Andor Borealis unit reshaped the laser beam from a Gaussian profile to a homogenous flat top. As a dichroic mirror, a CR-DFLY-DMQD-01 was used. Fluorescence light was spectrally filtered with an emission filter (TR-DFLY-F600-050) and imaged with a scientific complementary metal oxide semiconductor (sCMOS) camera (Andor Technologies) without further magnification, resulting in an effective pixel size of 100.5 nm.

Three-dimensional super-resolution imaging was performed by introducing astigmatism via a cylindrical lens in front of the camera^52^.

### Image acquisition

A detailed summary of imaging conditions for all DNA origami and cell experiments can be found in **Suppl. Table 3**.

### Image analysis

All raw fluorescence microscopy images were subjected to spot-finding and subsequent super-resolution reconstruction using the ‘Picasso’ software package^33^.

#### DNA origami analysis for 20-nm grids

Localized single-molecule data was drift corrected using redundant cross-correlation (RCC)^53^ and the adaptive intersection maximization-based (AIM) algorithm^54^, and, finally, resolution permitting, undrifted by interactively picked individual docking sites. DNA origami were visualized using the ‘PYMEVisualize’ software package^50^.

#### DNA origami analysis for kinetics quantification

Localized single-molecule data was subjected to drift correction and alignment using Picasso. For center docking site rounds (**Supp. Fig. 3c**), drift correction was performed with the AIM algorithm^54^; for frame rounds, drift correction was performed using RCC^53^ and interactively selected DNA origami structures. Localizations that appeared within a radius of twice the median localization precision and allowing 2 transient dark frames of misdetection were linked into ‘events’, taking into account that one blinking event can span multiple frames. Localizations were filtered for outliers using custom Python scripts. Single docking sites were identified by aligning the center docking site round to its corresponding frame round and interactively selecting circular ‘pick’ regions containing a single DNA origami structure. For the data in **Fig. 1d-h** and **Suppl. Fig. 3d-h**, each picked region was additionally filtered by performing a 2D Gaussian kernel density estimation using SciPy to extract localization events originating from the center docking site. Finally, Picasso was used to calculate kinetic properties for each docking site. Bulk kinetic analysis and plot generation were performed using custom Python scripts. For each probe, data was collected in n=3 independent experiments and in each experiment, values were averaged over ∼2000 origami). For the radar plot (**Fig. 1h**), each value was normalized to the maximum value in that category. Non-specific binding was evaluated qualitatively by visual inspection of the imaging data. Based on empirical assessment of image quality and background signal, three discrete scores were assigned to represent relative performance: 0.2 (poor), 0.6 (good), and 1 (excellent). Since lower values are more desirable for the localization precision, the values were inverted relative to those shown in the bar plot.

#### Microtubule data analysis

Localized single-molecule data was subjected to drift correction with gold nanoparticles as fiducials, filtering, and alignment using Picasso. The fitted Z-coordinate of each localization was determined based on the degree of astigmatism via a calibration file of a bead stack. Localizations outside of the ±400 nm Z-range were discarded. The microtubule segment shown was picked in Picasso and cross-sectional profiles were generated with custom Python scripts. Gold nanoparticles were excluded.

#### Telomere data analysis

Localized single-molecule data was subjected to drift correction with the AIM algorithm (cite) using Picasso. Localizations were filtered for outliers using custom Python scripts, then localizations that appeared within a radius of twice the median localization precision and allowing 2 transient dark frames of misdetection were linked into ‘events’, and finally, all three rounds of imaging for each nucleus were aligned. For rounds that showed telomere puncta, DBSCAN^55^ was performed with Picasso to determine cluster centers. Basic frame analysis was performed with custom scripts to only keep clusters with a mean frame value within 1500 frames of the expected average (half the acquisition time) and a standard deviation in frames greater than 6250 frames (to allow for some deviation from a true uniform distribution). Clusters with a convex hull area less than 3 px^2^ were considered out-of-plane telomeres and thus excluded from both signal and background calculations. Circular picked regions around the cluster centers were used as telomere signal. For all rounds, the nucleus was segmented by thresholding and manually selected circular picked regions evenly distributed throughout the segmented area were used as nuclear background. Localization density values were computed as total number of localizations in a picked region divided by the respective area. Data was collected from n=3 nuclei from three independent experiments with a total of N=43 telomere clusters and N=1566 nuclear background regions.

#### Myc-POT1 and TRF2 data analysis

After spot-fitting, a nuclear mask was first applied to both rounds to minimize the impact of noise on drift correction since these are low-abundance targets. Then, drift correction was performed using the AIM algorithm^54^, localizations were filtered for outliers using custom Python scripts, and finally, both rounds of imaging were aligned. Intensity profiles are calculated for a 30 nm-wide rectangular region centered at the straight, dotted line shown in zoomed-in views; the region was picked in Picasso and plots were generated using custom Python scripts. The two-color data was visualized using PYMEVisualize 23.5.17.0^50^.

#### Sec61β data analysis

The z-coordinate of each localization was determined based on the degree of astigmatism via a calibration file of a bead stack through fitting in Picasso and custom Python scripts. Localizations outside of the ±400 nm Z-range for each slice were discarded and filtered for outliers. Then, each slice was drift corrected using the AIM^54^ algorithm or gold nanoparticles as fiducials when possible and aligned to form one large stack. The localizations of the first two cycles are displayed after performing a DBSCAN^55^ to extract the densely-labeled ER structure and remove noise.

