## Supplementary_Information for "Fluorogenic speed-optimized DNA-PAINT probes enable super-resolution imaging of whole cells"

for

Supplementary Note 1: Performance metrics for DNA-PAINT probes.
Supplementary Note 2: Selection of laser intensities for FSP1 and FSP2.

Supplementary Figure 1: Factors impacting DNA-PAINT performance.
Supplementary Figure 2: Binding kinetics of FSPs under different laser intensities.
Supplementary Figure 3: Characterization of DNA-PAINT probes with DNA origami.
Supplementary Figure 4: DNA-PAINT images of microtubules in COS-7 cells with FSPs.
Supplementary Figure 5: DNA-PAINT images of telomeres in U-2 OS cell nuclei.

Supplementary Table 1: DNA-PAINT probe sequences.
Supplementary Table 2: Docking site and FLASH-PAINT sequences.
Supplementary Table 3: Imaging conditions.
Supplementary Table 4: DNA origami biotinylated staples. See Excel sheet.
Supplementary Table 5: DNA origami staples for 20-nm grids. See Excel sheet.
Supplementary Table 6: DNA origami staples for frames. See Excel sheet.

### Supplementary Note 1

**Performance metrics for DNA-PAINT probes**

The quality of a super-resolution DNA-PAINT image depends on:

1. the precision with which individual molecules can be localized, which depends on the number of photons per localized blinking event (in short: blink) and the background, and
2. the total number of 'blinks' that were localized, which is a measure of the sampling density in the image.

An equally important factor is the rate at which blinks are acquired since this determines how long a sample needs to be imaged to reach an acceptable sampling density of target molecules and hence describes the throughput of an imaging experiment.

While these factors can be extracted relatively easily from any DNA-PAINT data set, they strongly depend on the imaged sample, especially on the number of docking sites the used probes can bind to. For a less sample-dependent evaluation of DNA-PAINT probes, metrics describing the binding kinetics of the probes, in particular the association and dissociation rates of probes binding to a docking site, are better options.

We identified four metrics, described in more detail below, that we believe provide a concise yet comprehensive characterization of DNA-PAINT probes:

1. the bleaching-affected effective association rate (**Fig. 1d**),
2. the effective ON time (**Fig. 1e**),
3. the localization precision (**Fig. 1f**), and
4. a newly introduced Speed Value (**Fig. 1g**).

Bleaching-affected Effective Association Rate $k_{on}$

The association rate ($k_{on}$) of a probe is the rate at which a probe in solution binds to a docking site. It is measured in units of $M^{-1}s^{-1}$. When multiplied by the probe concentration, $k_{on}$ yields the binding rate ($s^{-1}$), describing the expected number of blinking events per second for a single docking site. The ideal probe has the highest possible association rate since this allows it to achieve a targeted blinking frequency at the lowest probe concentration and thus minimal background. In other words, for a given probe concentration, increasing the association rate decreases the time between binding events, allowing the required number of blinks to be obtained in a shorter amount of time (**Suppl. Fig. 1a**). The association rate is dependent on intrinsic properties of the sequence design[1,2] but can also be adjusted through buffer optimization as salt concentrations affect ssDNA binding affinity[3].

DNA-PAINT leverages a large, replenishable pool of probes in solution (**Supp. Fig 1c**, left column), making it less susceptible to photobleaching compared to SMLM approaches that rely on photoswitching of permanently bound fluorophores (e.g. STORM, PALM, FPALM, dSTORM)[4]. However, as the excitation volume increases, probe replenishment becomes less efficient. In widefield illumination, for example, probes must diffuse across long distances and may bleach in solution before ever reaching an intended target in the center of the field of view (**Supp. Fig. 1c**, top right). This results in a reduced number of observed blinks, resembling a lower effective association rate. The bleaching effect is particularly pronounced in thick or dense biological samples, where the crowded environment slows down diffusion, e.g., the cell nucleus. It also depends on the imaging conditions, including the size and thickness of the illuminated volume as well as the laser intensity. The ideal probe is robust against variations of these parameters.

Since photobleaching is associated with the excited state of the fluorophore, we reasoned that a probe will be resistant to bleaching while quenched. Therefore, FSPs should provide a more uniform availability of unbleached probes throughout the illuminated volume (**Supp. Fig. 1c**, bottom right).

Here, we measured the effective association rate as the inverse of the product of the probe concentration and the average time between probes being observed to be bound to our single-docking-site DNA origami test structures.

Effective ON time $1/k_{off}$

To minimize the number of camera frames, and therefore the data acquisition time needed to localize a specific number of blinking events with minimal interference between neighboring binding sites, each bound molecule should ideally be visible (i.e. be “ON”) for not more than a single camera frame. Similarly, there is no benefit of the ON time being substantially shorter than the exposure time for a single camera frame, since background photons will be accumulated over the full exposure time even if the ON time is shorter, which negatively impacts the localization precision (see below). The ideal ON time therefore matches the camera frame exposure time, or, in other words, the ideal effective dissociation rate $k_{off}$ matches the camera frame rate.

The ON time ($1/k_{off}$) primarily depends on the probe sequence design, buffer conditions (e.g., salinity), and temperature. In DNA-PAINT experiments, the measured ON time can additionally be influenced by photobleaching. High laser intensities can lead to premature bleaching of probes while they are still bound, resulting in an increased effective dissociation rate.

Here, we measured the effective ON time as the average time probes were observed to be bound to our DNA origami test structures.

Localization Precision

High localization precision is essential for achieving high-resolution SMLM images. This depends primarily on two factors: (i) the number of photons collected during a blinking event (i.e. the signal level), which appertains to the molecular brightness of the probe, and (ii) the background level, which can come from a variety of sources including autofluorescence, unbound probes in solution, and probes that are bound specifically or unspecifically nearby (including in planes above or below the focal plane). The localization precision of a single blinking event is known to be proportional to the full width at half maximum of a Gaussian curve fitted to the point spread function; thus, the combination of high signal level and low background level leads to the best localization precision (**Suppl. Fig. 1d**).

Speed Value

The throughput of a DNA-PAINT experiment depends on how rapidly each docking site within an ROI can be sampled by imager probes. While sampling can be increased by raising the probe concentration in solution, this has the disadvantage that higher probe concentrations increase the background, which negatively impacts localization precision. Conversely, lowering the probe concentration improves precision by reducing background, but decreases the sampling frequency and thus the overall imaging throughput.

The performance of a probe in terms of achievable localization precision and imaging speed therefore depends on the probe concentration. Notably, since both the background and the number of binding events scale in a good approximation proportionally with probe concentration, a concentration-independent quality metric for a probe can be obtained by dividing one by the other.

We therefore introduce a new metric, the ‘speed value’, as the number of binding events per second divided by the number of background photons per second. Probes that generate many blinks while maintaining a low background have a high speed value.

### Supplementary Note 2

To obtain an unbiased comparison between the FSP1 and FSP2 and the gold standards SP and FP, laser intensities for FSP1 and FSP2 had to be carefully selected to account for differences in the binding kinetics and photophysics of the probes.

When the laser intensity is too low, the observed kinetics can appear erratic and misleading because fluorophores are not sufficiently excited to be reliably detected in individual camera frames. At higher laser intensities, the fluorescence photon flux increases, improving both the detection efficiency and the localization precision per camera frame. However, excessively high laser intensities can also lead to non-linear bleaching effects, causing dyes to bleach prematurely while diffusing to the docking site and thereby reducing the average number of photons detected from each binding event.

This general behavior can be observed in **Suppl. Fig. 2**. For high intensities, measured effective $k_{on}$ values are low since a subset of probes bleach before they bind. Similarly, measured effective $k_{off}$ values are high since a fraction of molecules bleaches while bound, reducing the effective ON time.

At lower intensities, the same trend in measured $k_{on}$ and $k_{off}$ values can be observed, for a different reason: weak excitation results in low signal per camera frame, causing bound molecules to be ‘undetected,’ leading in turn to an artificially low detected number of binding events and reduced binding durations.

Because these artifacts arise from inefficient detection, we chose laser intensities approximately threefold higher than the range at which these detection-limit artifacts disappear, while still remaining low enough in laser intensity to avoid bleaching-dominated effects. While FSP1 performs optimally around 1-1.5 kW/cm$^2$ and shows increased bleaching above that range, FSP2 can withstand higher laser intensities of ~3-4 kW/cm$^2$ and be pushed to have a higher dissociation rate without sacrificing effective association rate to generate even faster binding kinetics.

For SP and FP, we chose intensity values of ~0.5 and ~4.1 kW/cm$^2$, respectively, similar to previously reported values[1,5]

### Supplementary Figure 1

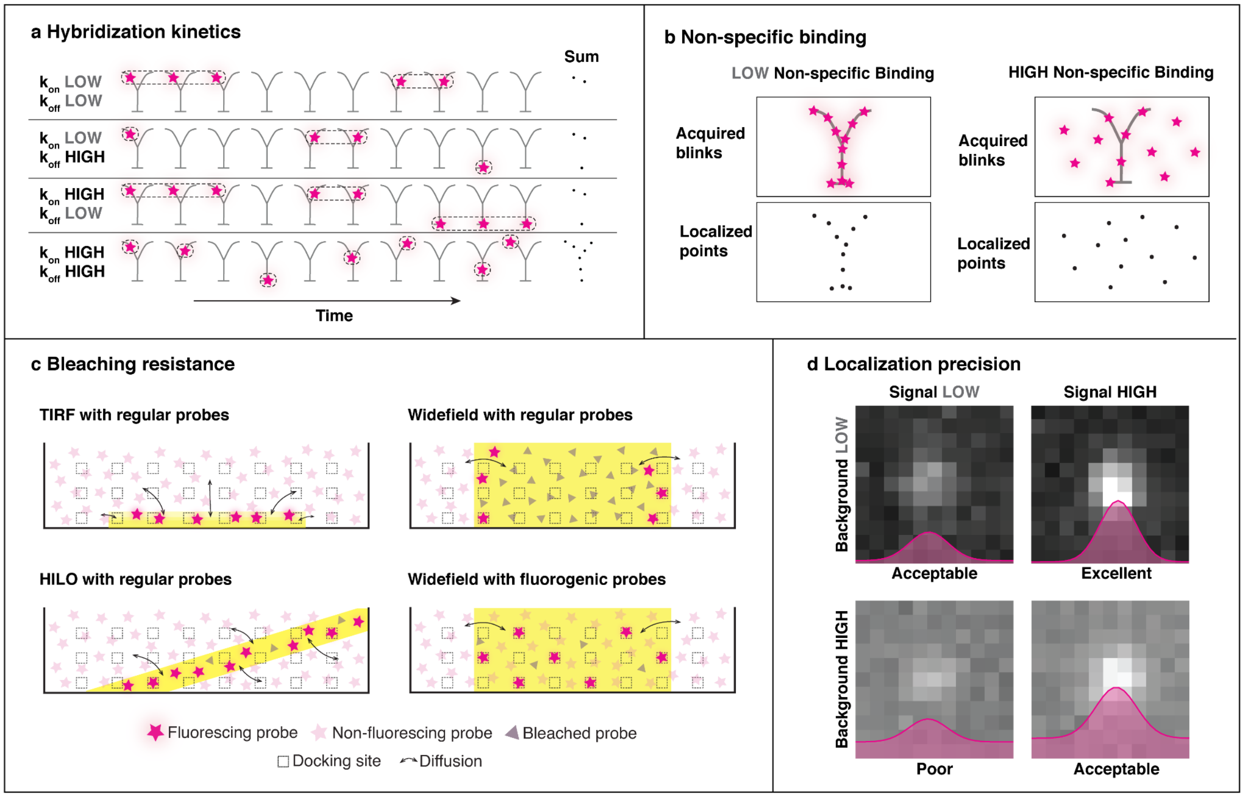


***Supplementary Figure 1. Factors impacting DNA-PAINT performance.*** *(a)* ***Hybridization kinetics****. Depiction of acquired 'blinks' in DNA-PAINT imaging over time for a Y-shaped ground truth structure under different* $k_{on}$ *and* $k_{off}$ *conditions. High* $k_{on}$ *and* $k_{off}$ *leads to fastest target sampling. Note this is not drawn to scale in time. (b)* ***Non-specific binding****. Depiction of acquired DNA-PAINT localizations over the same length of time when there is low or high non-specific binding (grey Y is ground truth structure). In the high non-specific binding case, the true structure is obfuscated. (c)* ***Bleaching resistance****. Depiction of bleaching effect under different types of illumination: TIRF, HILO, and widefield. (d)* ***Localization precision****. Depiction of single DNA-PAINT blinking event fitted with a Gaussian under different signal and background levels. The combination of high signal and low background yields the best localization precision.*

### Supplementary Figure 2

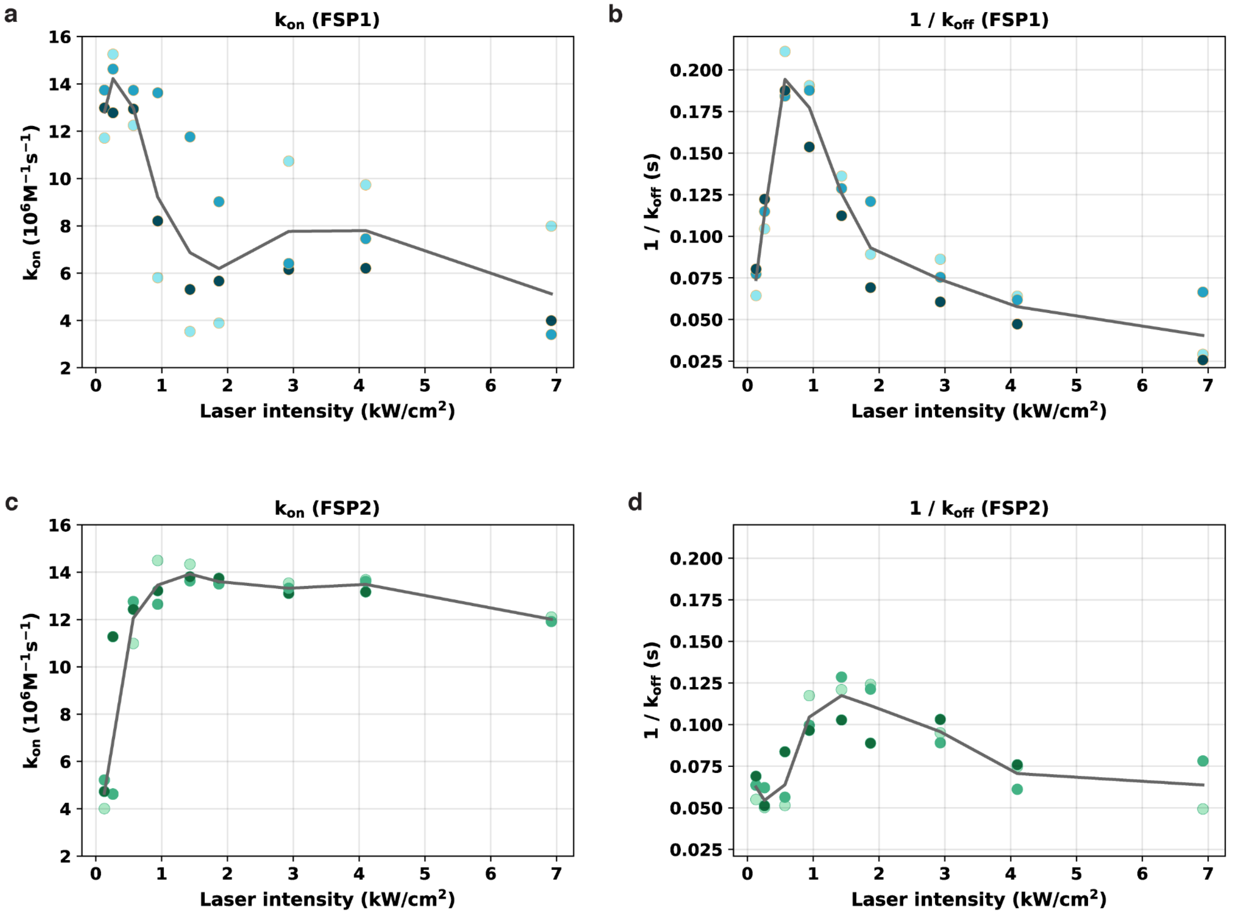


***Supplementary Figure 2. Binding kinetics of FSPs under different laser intensities.*** *The effective association rate ($k_{on}$) and ON time ($1/k_{off}$) measured with DNA origami (**Supp. Fig. 3c**) for laser intensities ranging from 0.13 to 6.92 kW/cm$^2$. (a-b) FSP1 binding kinetics ($k_{on}$ and $1/k_{off}$). (c-d) FSP2 binding kinetics ($k_{on}$ and $1/k_{off}$).*

### Supplementary Figure 3

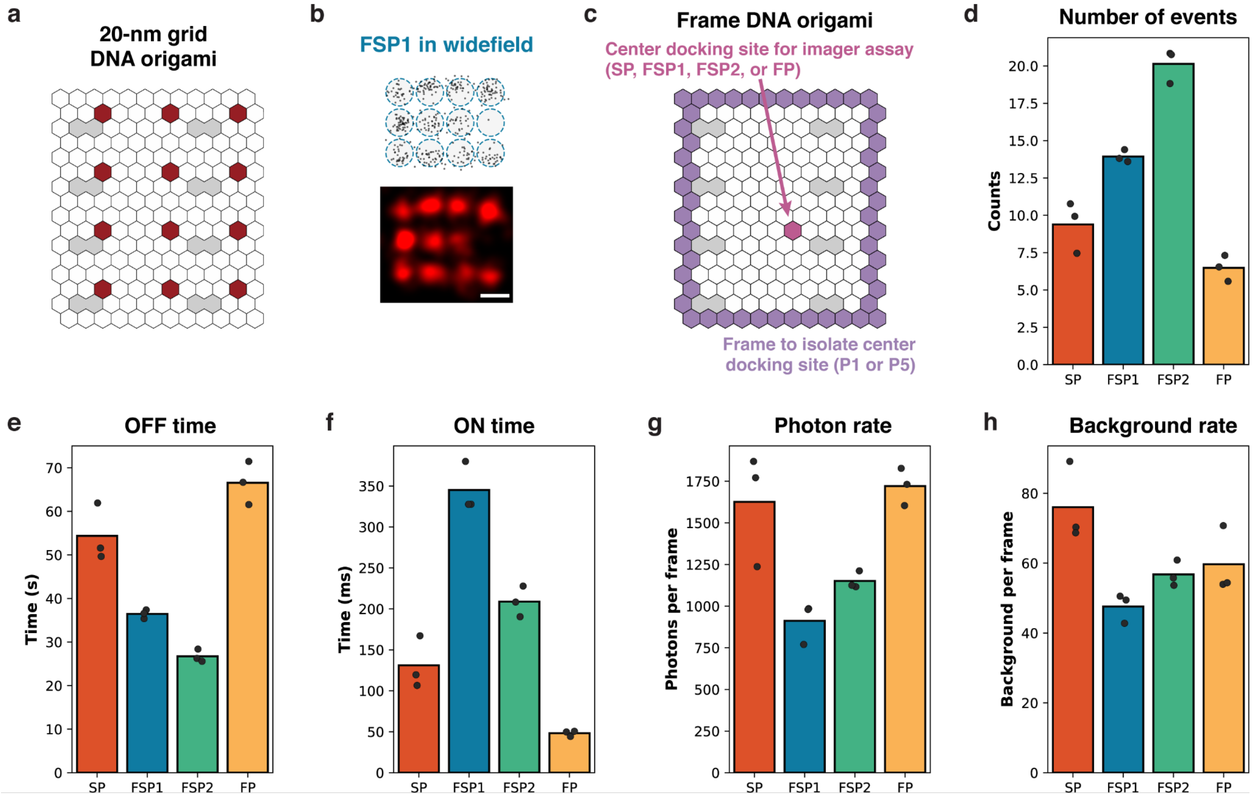


***Supplementary Figure 3. Characterization of DNA-PAINT probes with DNA origami.*** *(a) Schematic of 20-nm grid DNA origami structures. Red hexagons represent the positions of staples carrying the SP docking site, forming the grid structure. Grey hexagons represent the positions of staples modified with biotin to anchor the DNA origami to the coverglass surface. (b) DNA-PAINT imaging of 20-nm grid DNA origami demonstrating FSP1 can resolve the structure under widefield illumination. Scale bar: 20nm. (c) Schematic of DNA origami structure used for single-molecule characterization of probes. The center docking site (pink hexagon) from which the kinetics are calculated is either the SP or FP docking site. The purple hexagons carry the P1 or P5 docking site and form a 'frame' shape that is used to isolate individual center docking sites from each other and from any non-specific binding outside the origami. (d-h) Additional probe properties measured from DNA origami.*

### Supplementary Figure 4

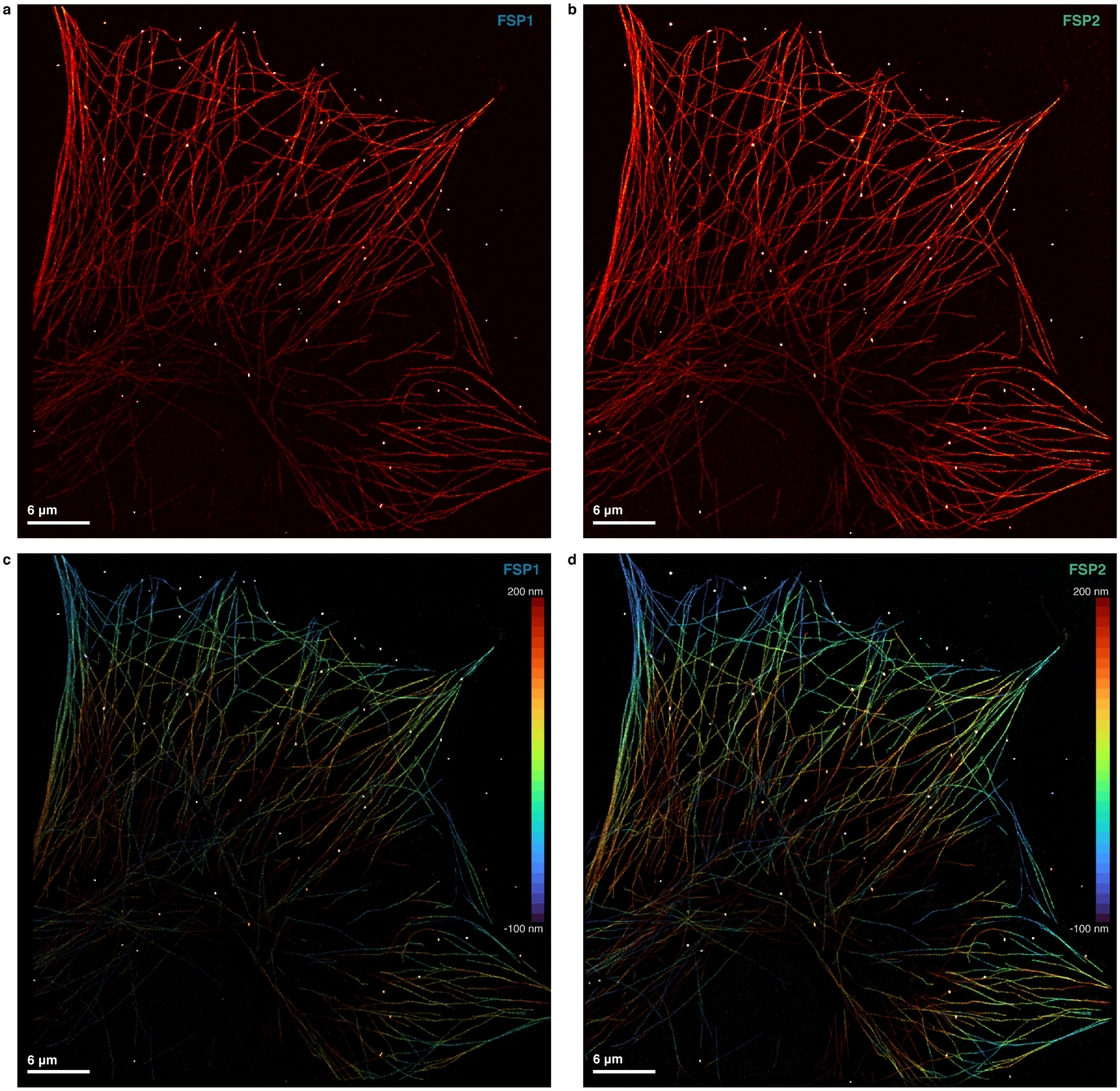


***Supplementary Figure 4. DNA-PAINT images of microtubules in COS-7 cells with FSPs.*** *(a) Full field-of-view image of microtubules imaged with FSP1 from Fig. 2a. (b) Full field-of-view image of microtubules imaged with FSP2 from Fig. 2a. (c-d) Images shown in a-b with pseudocoloring by Z-position over a 300 nm range.*

### Supplementary Figure 5

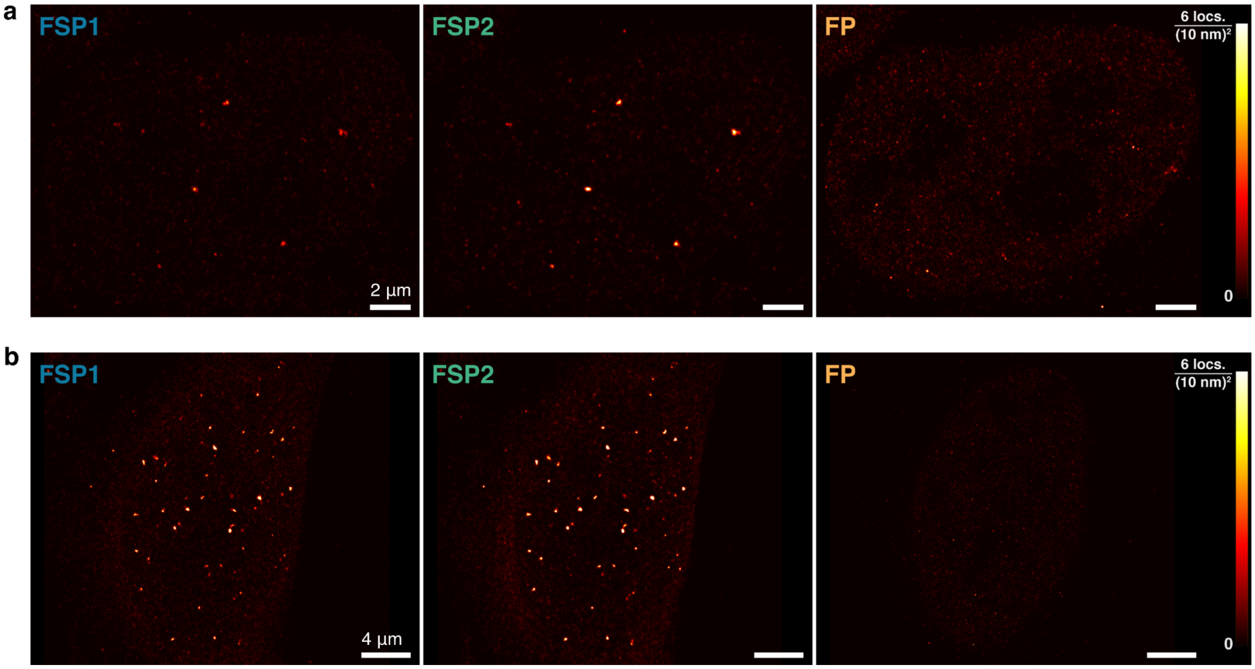


***Supplementary Figure 5. DNA-PAINT images of telomeres in U-2 OS cell nuclei.*** *(a-b) Three rounds of DNA-PAINT imaging of telomeres with FSP1, FSP2, and FP, each for a different U-2 OS cell nucleus for a total of n=3 independent repeats with Fig. 2d. Differences in total telomere puncta abundance are expected due to variation in hybridization efficiency and imaging plane.*

### Supplementary Table 1: DNA-PAINT probe sequences

| DNA-PAINT probe | 5' mod | Sequence | 3' mod |
|---|---|---|---|
| SP[1] | Cy3B | TGG TGG | |
| FP[5,6] | Cy3B | AAGAAGTAAAGGGAG | BHQ2 |
| P5 | | ATACATTGA | Cy3B |
| P1 | Cy3B | TAGATGTAT | |
| FSP1 | Cy3B - PEG9 | TGG TGG | PEG9 - IBRQ |
| FSP2 | Cy3B - PEG9 | TGG TGG | PEG9 - BHQ2 |

### Supplementary Table 2: Docking site and FLASH-PAINT sequences

| Docking sites | 5’ to 3’ sequence |
|---|---|
| SP docking site | ACCACCACCACC |
| FP docking site | ATTCCCCAAGTCGCTCC |
| P1 docking site | ATACATCTA |
| P5 docking site | TCAATGTAT |
| Telomere primary probe | CCTCGCTGAACCCCTTA AA CCCTAACCCTAACCCTAACCCTAA AA ACCACCACCACC |
| A5 docking site | TT TCAATGTATGGC |
| **FLASH-PAINT sequences** | **5’ to 3’ sequence** |
| A3-5xR2 | ACCACCACCACCACCACCA AA CGCTAATGAA |
| A5-FP2 | CCTCGCTGAACCCCTTA AA GCCATACATT |
| A5-5xR2 | ACCACCACCACCACCACCA AA GCCATACATT |
| A8-5xR2 | ACCACCACCACCACCACCA AA ACCCATTAAC |
| E3-R2 | TTCATTAGCG AA TGG |

### Supplementary Table 3: Imaging conditions

| Experiment | Imaging rounds | Buffer | Exposure & integration time | Laser intensity |
|---|---|---|---|---|
| Fig. 1c (TIRF) | SP (1 nM) | B+, 20mM $Na_2SO_3$, 2mM Trolox | 100 ms, 20,000 frames | 0.5 $kW/cm^2$ |
| Fig. 1c (widefield) | SP or FSP2 (8 nM) | B+, 20mM $Na_2SO_3$, 2mM Trolox | 100 ms, 20,000 frames | SP - 0.4 $kW/cm^2$<br>FSP2 - 3.5 $kW/cm^2$ |
| Fig. 1d-h, Suppl. Fig. 2, 3d-h | Center docking site - SP, FSP1, FSP2, or FP (5 nM) | B+ | 20 ms, 25,000 frames | SP - 0.5 $kW/cm^2$<br>FSP1 - 1.5 $kW/cm^2$<br>FSP2 - 4.1 $kW/cm^2$<br>FP - 4.1 $kW/cm^2$ |
| | Frame - P1 or P5 (5 nM) | B+ | 50 ms, 5,000 frames | 0.5 $kW/cm^2$ |
| Fig. 2a-c, Suppl. Fig. 4 | FSP1 (400 pM), A3-5xR2 (20 nM) | C, 20mM $Na_2SO_3$, 1mM Trolox | 100 ms, 25,000 frames | 1.9 $kW/cm^2$ |
| | FSP2 (100 pM), A3-5xR2 (20 nM) | | 50 ms, 50,000 frames | 4.1 $kW/cm^2$ |
| Fig. 2d-f, Suppl. Fig. 5 | FSP1, FSP2, and FP (1 nM) | C | 20 ms, 25,000 frames | 1.3 $kW/cm^2$ |
| Fig. 2g | Myc-POT1 - FSP1 (5 nM), A3-5xR2 (20n M) | C | 20 ms, 40,000 frames | 1.3 $kW/cm^2$ |
| | TRF2 - FSP1 (5 nM), A8-5xR2 (20 nM) | | | |
| Fig. 3 | FSP2 (500 pM) | C, 20mM $Na_2SO_3$, 1mM Trolox | 50 ms, 30,000 frames per Z-step, 21 slices per stack, six total stack acquired | 4.1 $kW/cm^2$ |
| Suppl. Fig. 3b | FSP1 (5 nM) | B+, 20mM $Na_2SO_3$, 1mM Trolox | 120 ms, 16,667 frames | 1.9 $kW/cm^2$ |

### Supplementary Table 4 - DNA origami biotinylated staples

See file: SupplementaryTable4_biotinylated-staples.xslx

### Supplementary Table 5 - DNA origami staples for 20-nm grids

See file: SupplementaryTable5_biotinylated-staples.xslx

### Supplementary Table 6 - DNA origami staples for Frames

See file: SupplementaryTable6_frame-staples.xslx
