## Supplementary_Table_4 for "Fluorogenic speed-optimized DNA-PAINT probes enable super-resolution imaging of whole cells"

| DNA origami biotinylated staples | | | | |
|---|---|---|---|---|
| **Plate name** | **Plate position** | **Oligo name** | **Sequence** | **5' mod** |
| BIOTIN | C2 | 18[63]20[56]BIOTIN | ATTAAGTTTACCGAGCTCGAATTCGGGAAACCTGTCGTGC | 5' Biotin |
| BIOTIN | C3 | 4[63]6[56]BIOTIN | ATAAGGGAACCGGATATTCATTACGTCAGGACGTTGGGAA | 5' Biotin |
| BIOTIN | C9 | 18[127]20[120]BIOTIN | GCGATCGGCAATTCCACACAACAGGTGCCTAATGAGTG | 5' Biotin |
| BIOTIN | C10 | 4[127]6[120]BIOTIN | TTGTGTCGTGACGAGAAACACCAAATTTCAACTTTAAT | 5' Biotin |
| BIOTIN | G2 | 18[191]20[184]BIOTIN | ATTCATTTTTGTTTGGATTATACTAAGAAACCACCAGAAG | 5' Biotin |
| BIOTIN | G3 | 4[191]6[184]BIOTIN | CACCCTCAGAAACCATCGATAGCATTGAGCCATTTGGGAA | 5' Biotin |
| BIOTIN | G9 | 18[255]20[248]BIOTIN | AACAATAACGTAAAACAGAAATAAAAATCCTTTGCCCGAA | 5' Biotin |
| BIOTIN | G10 | 4[255]6[248]BIOTIN | AGCCACCACTGTAGCGCGTTTTCAAGGGAGGGAAGGTAAA | 5' Biotin |
