## Supplementary_Table_5 for "Fluorogenic speed-optimized DNA-PAINT probes enable super-resolution imaging of whole cells"

| 20-nm grid staples | | | |
|---|---|---|---|
| **Plate name** | **Plate position** | **Oligo name** | **Sequence** |
| BLK_1 | A1 | 21[32]23[31]BLK | TTTTCACTCAAAGGGCGAAAAACCATCACC |
| BLK_1 | A2 | 19[32]21[31]BLK | GTCGACTTCGGCCAACGCGCGGGGTTTTTC |
| BLK_1 | A3 | 17[32]19[31]BLK | TGCATCTTTCCCAGTCACGACGGCCTGCAG |
| BLK_1 | A4 | 15[32]17[31]BLK | TAATCAGCGGATTGACCGTAATCGTAACCG |
| BLK_1 | A5 | 13[32]15[31]BLK | AACGCAAAATCGATGAACGGTACCGGTTGA |
| BLK_1 | A6 | 11[32]13[31]BLK | AACAGTTTTGTACCAAAAACATTTTATTTC |
| BLK_1 | A7 | 9[32]11[31]BLK | TTTACCCCAACATGTTTTAAATTTCCATAT |
| BLK_1 | A8 | 7[32]9[31]BLK | TTTAGGACAAATGCTTTAAACAATCAGGTC |
| BLK_1 | A9 | 5[32]7[31]BLK | CATCAAGTAAAACGAACTAACGAGTTGAGA |
| BLK_1 | A10 | 3[32]5[31]BLK | AATACGTTTGAAAGAGGACAGACTGACCTT |
| BLK_1 | A11 | 1[32]3[31]BLK | AGGCTCCAGAGGCTTTGAGGACACGGGTAA |
| BLK_1 | A12 | 0[47]1[31]BLK | AGAAAGGAACAACTAAAGGAATTCAAAAAAA |
| BLK_1 | B1 | 23[32]22[48]BLK | CAAATCAAGTTTTTTGGGGTCGAAACGTGGA |
| BLK_1 | B2 | 22[47]20[48]BLK | CTCCAACGCAGTGAGACGGGCAACCAGCTGCA |
| BLK_1 | B3 | 20[47]18[48] - SP docking strand | TTAATGAACTAGAGGATCCCCGGGGGGTAACG AAACCACCACCACCACCACCA |
| BLK_1 | B4 | 18[47]16[48]BLK | CCAGGGTTGCCAGTTTGAGGGGACCCGTGGGA |
| BLK_1 | B5 | 16[47]14[48]BLK | ACAAACGGAAAAGCCCCAAAAACACTGGAGCA |
| BLK_1 | B6 | 14[47]12[48]BLK | AACAAGAGGGATAAAAATTTTTAGCATAAAGC |
| BLK_1 | B7 | 12[47]10[48] - SP docking strand | TAAATCGGGATTCCCAATTCTGCGATATAATG AAACCACCACCACCACCACCA |
| BLK_1 | B8 | 10[47]8[48]BLK | CTGTAGCTTGACTATTATAGTCAGTTCATTGA |
| BLK_1 | B9 | 8[47]6[48]BLK | ATCCCCCTATACCACATTCAACTAGAAAAATC |
| BLK_1 | B10 | 6[47]4[48]BLK | TACGTTAAAGTAATCTTGACAAGAACCGAACT |
| BLK_1 | B11 | 4[47]2[48] - SP docking strand | GACCAACTAATGCCACTACGAAGGGGGTAGCA AAACCACCACCACCACCACCA |
| BLK_1 | B12 | 2[47]0[48]BLK | ACGGCTACAAAAGGAGCCTTTAATGTGAGAAT |
| BLK_1 | C1 | 21[56]23[63]BLK | AGCTGATTGCCCTTCAGAGTCCACTATTAAAGGGTGCCGT |
| BIOTIN | C2 | | |
| BIOTIN | C3 | | |
| BLK_1 | C4 | 15[64]18[64]BLK | GTATAAGCCAACCCGTCGGATTCTGACGACAGTATCGGCCGCAAGGCG |
| BLK_1 | C5 | 13[64]15[63]BLK | TATATTTTGTCATTGCCTGAGAGTGGAAGATT |
| BLK_1 | C6 | 11[64]13[63]BLK | GATTTAGTCAATAAAGCCTCAGAGAACCCTCA |
| BLK_1 | C7 | 9[64]11[63]BLK | CGGATTGCAGAGCTTAATTGCTGAAACGAGTA |
| BLK_1 | C8 | 7[56]9[63]BLK | ATGCAGATACATAACGGGAATCGTCATAAATAAAGCAAAG |
| BIOTIN | C9 | | |
| BIOTIN | C10 | | |
| BLK_1 | C11 | 1[64]4[64]BLK | TTTATCAGGACAGCATCGGAACGACACCAACCTAAAACGAGGTCAATC |
| BLK_1 | C12 | 0[79]1[63]BLK | ACAACTTTCAACAGTTTCAGCGGATGTATCGG |
| BLK_1 | D1 | 23[64]22[80]BLK | AAAGCACTAAATCGGAACCCTAATCCAGTT |
| BLK_1 | D2 | 22[79]20[80]BLK | TGGAACAACCGCCTGGCCCTGAGGCCCGCT |
| BLK_1 | D3 | 20[79]18[80]BLK | TTCCAGTCGTAATCATGGTCATAAAAGGGG |
| BLK_1 | D4 | 18[79]16[80]BLK | GATGTGCTTCAGGAAGATCGCACAATGTGA |
| BLK_1 | D5 | 16[79]14[80]BLK | GCGAGTAAAAATATTTAAATTGTTACAAAG |
| BLK_1 | D6 | 14[79]12[80]BLK | GCTATCAGAAATGCAATGCCTGAATTAGCA |
| BLK_1 | D7 | 12[79]10[80]BLK | AAATTAAGTTGACCATTAGATACTTTTGCG |
| BLK_1 | D8 | 10[79]8[80]BLK | GATGGCTTATCAAAAAGATTAAGAGCGTCC |
| BLK_1 | D9 | 8[79]6[80]BLK | AATACTGCCCAAAAGGAATTACGTGGCTCA |
| BLK_1 | D10 | 6[79]4[80]BLK | TTATACCACCAAATCAACGTAACGAACGAG |
| BLK_1 | D11 | 4[79]2[80]BLK | GCGCAGACAAGAGGCAAAAGAATCCCTCAG |
| BLK_1 | D12 | 2[79]0[80]BLK | CAGCGAAACTTGCTTTCGAGGTGTTGCTAA |
| BLK_1 | E1 | 21[96]23[95]BLK | AGCAAGCGTAGGGTTGAGTGTTGTAGGGAGCC |
| BLK_1 | E2 | 19[96]21[95]BLK | CTGTGTGATTGCGTTGCGCTCACTAGAGTTGC |
| BLK_1 | E3 | 17[96]19[95]BLK | GCTTTCCGATTACGCCAGCTGGCGGCTGTTTC |
| BLK_1 | E4 | 15[96]17[95]BLK | ATATTTTGGCTTTCATCAACATTATCCAGCCA |
| BLK_1 | E5 | 13[96]15[95]BLK | TAGGTAAACTATTTTTGAGAGATCAAACGTTA |
| BLK_1 | E6 | 11[96]13[95]BLK | AATGGTCAACAGGCAAGGCAAAGAGTAATGTG |
| BLK_1 | E7 | 9[96]11[95]BLK | CGAAAGACTTTGATAAGAGGTCATATTTCGCA |
| BLK_1 | E8 | 7[96]9[95]BLK | TAAGAGCAAATGTTTAGACTGGATAGGAAGCC |
| BLK_1 | E9 | 5[96]7[95]BLK | TCATTCAGATGCGATTTTAAGAACAGGCATAG |
| BLK_1 | E10 | 3[96]5[95]BLK | ACACTCATCCATGTTACTTAGCCGAAAGCTGC |
| BLK_1 | E11 | 1[96]3[95]BLK | AAACAGCTTTTTGCGGGATCGTCAACACTAAA |
| BLK_1 | E12 | 0[111]1[95]BLK | TAAATGAATTTTCTGTATGGGATTAATTTCTT |
| BLK_1 | F1 | 23[96]22[112]BLK | CCCGATTTAGAGCTTGACGGGGAAAAAGAATA |
| BLK_1 | F2 | 22[111]20[112]BLK | GCCCGAGAGTCCACGCTGGTTTGCAGCTAACT |
| BLK_1 | F3 | 20[111]18[112] - SP docking strand | CACATTAAAATTGTTATCCGCTCATGCGGGCC AAACCACCACCACCACCACCA |
| BLK_1 | F4 | 18[111]16[112]BLK | TCTTCGCTGCACCGCTTCTGGTGCGGCCTTCC |
| BLK_1 | F5 | 16[111]14[112]BLK | TGTAGCCATTAAAATTCGCATTAAATGCCGGA |
| BLK_1 | F6 | 14[111]12[112]BLK | GAGGGTAGGATTCAAAAGGGTGAGACATCCAA |
| BLK_1 | F7 | 12[111]10[112] - SP docking strand | TAAATCATATAACCTGTTTAGCTAACCTTTAA AAACCACCACCACCACCACCA |
| BLK_1 | F8 | 10[111]8[112]BLK | TTGCTCCTTTCAAATATCGCGTTTGAGGGGGT |
| BLK_1 | F9 | 8[111]6[112]BLK | AATAGTAAACACTATCATAACCCTCATTGTGA |
| BLK_1 | F10 | 6[111]4[112]BLK | ATTACCTTTGAATAAGGCTTGCCCAAATCCGC |
| BLK_1 | F11 | 4[111]2[112] - SP docking strand | GACCTGCTCTTTGACCCCCAGCGAGGGAGTTA AAACCACCACCACCACCACCA |
| BLK_1 | F12 | 2[111]0[112]BLK | AAGGCCGCTGATACCGATAGTTGCGACGTTAG |
| BLK_1 | G1 | 21[120]23[127]BLK | CCCAGCAGGCGAAAAATCCCTTATAAATCAAGCCGGCG |
| BIOTIN | G2 | | |
| BIOTIN | G3 | | |
| BLK_1 | G4 | 15[128]18[128]BLK | TAAATCAAAATAATTCGCGTCTCGGAAACCAGGCAAAGGGAAGG |
| BLK_1 | G5 | 13[128]15[127]BLK | GAGACAGCTAGCTGATAAATTAATTTTTGT |
| BLK_1 | G6 | 11[128]13[127]BLK | TTTGGGGATAGTAGTAGCATTAAAAGGCCG |
| BLK_1 | G7 | 9[128]11[127]BLK | GCTTCAATCAGGATTAGAGAGTTATTTTCA |
| BLK_1 | G8 | 7[120]9[127]BLK | CGTTTACCAGACGACAAAGAAGTTTTGCCATAATTCGA |
| BIOTIN | G9 | | |
| BIOTIN | G10 | | |
| BLK_1 | G11 | 1[128]4[128]BLK | TGACAACTCGCTGAGGCTTGCATTATACCAAGCGCGATGATAAA |
| BLK_1 | G12 | 0[143]1[127]BLK | TCTAAAGTTTTGTCGTCTTTCCAGCCGACAA |
| BLK_1 | H1 | 21[160]22[144]BLK | TCAATATCGAACCTCAAATATCAATTCCGAAA |
| BLK_1 | H2 | 19[160]20[144]BLK | GCAATTCACATATTCCTGATTATCAAAGTGTA |
| BLK_1 | H3 | 17[160]18[144]BLK | AGAAAACAAAGAAGATGATGAAACAGGCTGCG |
| BLK_1 | H4 | 15[160]16[144]BLK | ATCGCAAGTATGTAAATGCTGATGATAGGAAC |

| | | | |
|---|---|---|---|
| BLK_1 | H5 | 13[160]14[144]BLK | GTAATAAGTTAGGCAGAGGCATTTATGATATT |
| BLK_1 | H6 | 11[160]12[144]BLK | CCAATAGCTCATCGTAGGAATCATGGCATCAA |
| BLK_1 | H7 | 9[160]10[144]BLK | AGAGAGAAAAAAATGAAAATAGCAAGCAAACT |
| BLK_1 | H8 | 7[160]8[144]BLK | TTATTACGAAGAACTGGCATGATTGCGAGAGG |
| BLK_1 | H9 | 5[160]6[144]BLK | GCAAGGCCTCACCAGTAGCACCATGGGCTTGA |
| BLK_1 | H10 | 3[160]4[144]BLK | TTGACAGGCCACCACCAGAGCCGCGATTTGTA |
| BLK_1 | H11 | 1[160]2[144]BLK | TTAGGATTGGCTGAGACTCCTCAATAACCGAT |
| BLK_1 | H12 | 0[175]0[144]BLK | TCCACAGACAGCCCTCATAGTTAGCGTAACGA |
| BLK_2 | A1 | 23[128]23[159]BLK | AACGTGGCGAGAAAGGAAGGGAAACCAGTAA |
| BLK_2 | A2 | 22[143]21[159]BLK | TCGGCAAATCCTGTTTGATGGTGGACCCTCAA |
| BLK_2 | A3 | 20[143]19[159]BLK | AAGCCTGGTACGAGCCGGAAGCATAGATGATG |
| BLK_2 | A4 | 18[143]17[159]BLK | CAACTGTTGCGCCATTCGCCATTCAAACATCA |
| BLK_2 | A5 | 16[143]15[159]BLK | GCCATCAAGCTCATTTTTTAACCACAAATCCA |
| BLK_2 | A6 | 14[143]13[159]BLK | CAACCGTTTCAAATCACCATCAATTCGAGCCA |
| BLK_2 | A7 | 12[143]11[159]BLK | TTCTACTACGCGAGCTGAAAAGGTTACCGCGC |
| BLK_2 | A8 | 10[143]9[159]BLK | CCAACAGGAGCGAACCAGACCGGAGCCTTTAC |
| BLK_2 | A9 | 8[143]7[159]BLK | CTTTTGCAGATAAAAACCAAAATAAAGACTCC |
| BLK_2 | A10 | 6[143]5[159]BLK | GATGGTTTGAACGAGTAGTAAATTTACCATTA |
| BLK_2 | A11 | 4[143]3[159]BLK | TCATCGCCAACAAAGTACAACGGACGCCAGCA |
| BLK_2 | A12 | 2[143]1[159]BLK | ATATTCGGAACCATCGCCCACGCAGAGAAGGA |
| BLK_2 | B1 | 23[160]22[176]BLK | TAAAAGGGACATTCTGGCCAACAAAGCATC |
| BLK_2 | B2 | 22[175]20[176]BLK | ACCTTGCTTGGTCAGTTGGCAAAGAGCGGA |
| BLK_2 | B3 | 20[175]18[176] - SP docking strand | ATTATCATTCAATATAATCCTGACAATTAC AAACCACCACCACCACCACCA |
| BLK_2 | B4 | 18[175]16[176]BLK | CTGAGCAAAAATTAATTACATTTTGGGTTA |
| BLK_2 | B5 | 16[175]14[176]BLK | TATAACTAACAAAGAACGCGAGAACGCCAA |
| BLK_2 | B6 | 14[175]12[176]BLK | CATGTAATAGAATATAAAGTACCAAGCCGT |
| BLK_2 | B7 | 12[175]10[176] - SP docking strand | TTTTATTTAAGCAAATCAGATATTTTTTGT AAACCACCACCACCACCACCA |
| BLK_2 | B8 | 10[175]8[176]BLK | TTAACGTCTAACATAAAAACAGGTAACGGA |
| BLK_2 | B9 | 8[175]6[176]BLK | ATACCCAACAGTATGTTAGCAAATTAGAGC |
| BLK_2 | B10 | 6[175]4[176]BLK | CAGCAAAAGGAAACGTCACCAATGAGCCGC |
| BLK_2 | B11 | 4[175]2[176] - SP docking strand | CACCAGAAAGGTTGAGGCAGGTCATGAAAG AAACCACCACCACCACCACCA |
| BLK_2 | B12 | 2[175]0[176]BLK | TATTAAGAAGCGGGGTTTTGCTCGTAGCAT |
| BLK_2 | C1 | 21[184]23[191]BLK | TCAACAGTTGAAAGGAGCAAATGAAAAATCTAGAGATAGA |
| BIOTIN | C2 | | |
| BIOTIN | C3 | | |
| BLK_2 | C4 | 15[192]18[192]BLK | TCAAATATAACCTCCGGCTTAGGTAACAATTTCATTTGAAGGCGAATT |
| BLK_2 | C5 | 13[192]15[191]BLK | GTAAAGTAATCGCCATATTTAACAAAACTTTT |
| BLK_2 | C6 | 11[192]13[191]BLK | TATCCGGTCTCATCGAGAACAAGCGACAAAAG |
| BLK_2 | C7 | 9[192]11[191]BLK | TTAGACGGCCAAATAAGAAACGATAGAAGGCT |
| BLK_2 | C8 | 7[184]9[191]BLK | CGTAGAAAATACATACCGAGGAAACGCAATAAGAAGCGCA |
| BIOTIN | C9 | | |
| BIOTIN | C10 | | |
| BLK_2 | C11 | 1[192]4[192]BLK | GCGGATAACCTATTATTCTGAAACAGACGATTGGCCTTGAAGAGCCAC |
| BLK_2 | C12 | 0[207]1[191]BLK | TCACCAGTACAAACTACAACGCCTAGTACCAG |
| BLK_2 | D1 | 23[192]22[208]BLK | ACCCTTCTGACCTGAAAGCGTAAGACGCTGAG |
| BLK_2 | D2 | 22[207]20[208]BLK | AGCCAGCAATTGAGGAAGGTTATCATCATTTT |
| BLK_2 | D3 | 20[207]18[208]BLK | GCGGAACATCTGAATAATGGAAGGTACAAAAT |
| BLK_2 | D4 | 18[207]16[208]BLK | CGCGCAGATTACCTTTTTTAATGGGAGAGACT |
| BLK_2 | D5 | 16[207]14[208]BLK | ACCTTTTTATTTTAGTTAATTTCATAGGGCTT |
| BLK_2 | D6 | 14[207]12[208]BLK | AATTGAGAATTCTGTCCAGACGACTAAACCAA |
| BLK_2 | D7 | 12[207]10[208]BLK | GTACCGCAATTCTAAGAACGCGAGTATTATTT |
| BLK_2 | D8 | 10[207]8[208]BLK | ATCCCAATGAGAATTAACTGAACAGTTACCAG |
| BLK_2 | D9 | 8[207]6[208]BLK | AAGGAAACATAAAGGTGGCAACATTATCACCG |
| BLK_2 | D10 | 6[207]4[208]BLK | TCACCGACGCACCGTAATCAGTAGCAGAACCG |
| BLK_2 | D11 | 4[207]2[208]BLK | CCACCCTCTATTCACAAACAAATACCTGCCTA |
| BLK_2 | D12 | 2[207]0[208]BLK | TTTCGGAAGTGCCGTCGAGAGGGTGAGTTTCG |
| BLK_2 | E1 | 21[224]23[223]BLK | CTTTAGGGCCTGCAACAGTGCCAATACGTG |
| BLK_2 | E2 | 19[224]21[223]BLK | CTACCATAGTTTGAGTAACATTTAAAATAT |
| BLK_2 | E3 | 17[224]19[223]BLK | CATAAATCTTTGAATACCAAGTGTTAGAAC |
| BLK_2 | E4 | 15[224]17[223]BLK | CCTAAATCAAAATCATAGGTCTAAACAGTA |
| BLK_2 | E5 | 13[224]15[223]BLK | ACAACATGCCAACGCTCAACAGTCTTCTGA |
| BLK_2 | E6 | 11[224]13[223]BLK | GCGAACCTCCAAGAACGGGTATGACAATAA |
| BLK_2 | E7 | 9[224]11[223]BLK | AAAGTCACAAAATAAACAGCCAGCGTTTTA |
| BLK_2 | E8 | 7[224]9[223]BLK | AACGCAAAGATAGCCGAACAAACCCTGAAC |
| BLK_2 | E9 | 5[224]7[223]BLK | TCAAGTTTCATTAAAGGTGAATATAAAAGA |
| BLK_2 | E10 | 3[224]5[223]BLK | TTAAAGCCAGAGCCGCCACCCTCGACAGAA |
| BLK_2 | E11 | 1[224]3[223]BLK | GTATAGCAAACAGTTAATGCCCAATCCTCA |
| BLK_2 | E12 | 0[239]1[223]BLK | AGGAACCCATGTACCGTAACACTTGATATAA |
| BLK_2 | F1 | 23[224]22[240]BLK | GCACAGACAATATTTTTGAATGGGGTCAGTA |
| BLK_2 | F2 | 22[239]20[240]BLK | TTAACACCAGCACTAACAACTAATCGTTATTA |
| BLK_2 | F3 | 20[239]18[240] - SP docking strand | ATTTTAAAATCAAAATTATTTGCACGGATTCG AAACCACCACCACCACCACCA |
| BLK_2 | F4 | 18[239]16[240]BLK | CCTGATTGCAATATATGTGAGTGATCAATAGT |
| BLK_2 | F5 | 16[239]14[240]BLK | GAATTTATTTAATGGTTTGAAATATTCTTACC |
| BLK_2 | F6 | 14[239]12[240]BLK | AGTATAAAGTTCAGCTAATGCAGATGTCTTTC |
| BLK_2 | F7 | 12[239]10[240] - SP docking strand | CTTATCATTCCCGACTTGCGGGAGCCTAATTT AAACCACCACCACCACCACCA |
| BLK_2 | F8 | 10[239]8[240]BLK | GCCAGTTAGAGGGTAATTGAGCGCTTTAAGAA |
| BLK_2 | F9 | 8[239]6[240]BLK | AAGTAAGCAGACACCACGGAATAATATTGACG |
| BLK_2 | F10 | 6[239]4[240]BLK | GAAATTATTGCCTTTAGCGTCAGACCGGAACC |
| BLK_2 | F11 | 4[239]2[240] - SP docking strand | GCCTCCCTCAGAATGGAAAGCGCAGTAACAGT AAACCACCACCACCACCACCA |
| BLK_2 | F12 | 2[239]0[240]BLK | GCCCGTATCCGGAATAGGTGTATCAGCCCAAT |
| BLK_2 | G1 | 21[248]23[255]BLK | AGATTAGAGCCGTCAAAAAACAGAGGTGAGGCCTATTAGT |
| BIOTIN | G2 | | |
| BIOTIN | G3 | | |
| BLK_2 | G4 | 15[256]18[256]BLK | GTGATAAAAAGACGCTGAGAAGAGATAACCTTGCTTCTGTTCGGGAGA |
| BLK_2 | G5 | 13[256]15[255]BLK | GTTTATCAATATGCGTTATACAAACCGACCGT |
| BLK_2 | G6 | 11[256]13[255]BLK | GCCTTAAACCAATCAATAATCGGCACGCGCCT |
| BLK_2 | G7 | 9[256]11[255]BLK | GAGAGATAGAGCGTCTTTCCAGAGGTTTTGAA |
| BLK_2 | G8 | 7[248]9[255]BLK | GTTTATTTTGTCACAATCTTACCGAAGCCCTTTAATATCA |
| BIOTIN | G9 | | |
| BIOTIN | G10 | | |
| BLK_2 | G11 | 1[256]4[256]BLK | CAGGAGGTGGGGTCAGTGCCTTGAGTCTCTGAATTTACCGGGAACCAG |

| | | | |
|---|---|---|---|
| BLK_2 | G12 | 0[271]1[255]BLK | CCACCCTCATTTTCAGGGATAGCAACCGTACT |
| BLK_2 | H1 | 23[256]22[272]BLK | CTTTAATGCGCGAACTGATAGCCCCACCAG |
| BLK_2 | H2 | 22[271]20[272]BLK | CAGAAGATTAGATAATACATTTGTCGACAA |
| BLK_2 | H3 | 20[271]18[272]BLK | CTCGTATTAGAAATTGCGTAGATACAGTAC |
| BLK_2 | H4 | 18[271]16[272]BLK | CTTTTACAAAATCGTCGCTATTAGCGATAG |
| BLK_2 | H5 | 16[271]14[272]BLK | CTTAGATTTAAGGCGTTAAATAAAGCCTGT |
| BLK_2 | H6 | 14[271]12[272]BLK | TTAGTATCACAATAGATAAGTCCACGAGCA |
| BLK_2 | H7 | 12[271]10[272]BLK | TGTAGAAATCAAGATTAGTTGCTCTTACCA |
| BLK_2 | H8 | 10[271]8[272]BLK | ACGCTAACACCCACAAGAATTGAAAATAGC |
| BLK_2 | H9 | 8[271]6[272]BLK | AATAGCTATCAATAGAAAATTCAACATTCA |
| BLK_2 | H10 | 6[271]4[272]BLK | ACCGATTGTCGGCATTTTCGGTCATAATCA |
| BLK_2 | H11 | 4[271]2[272]BLK | AAATCACCTTCCAGTAAGCGTCAGTAATAA |
| BLK_2 | H12 | 2[271]0[272]BLK | GTTTTAACTTAGTACCGCCACCCAGAGCCA |
